# Dynamics of the bacterial community associated with *Phaeodactylum tricornutum* cultures

**DOI:** 10.1101/077768

**Authors:** Fiona Wanjiku Moejes, Ovidiu Popa, Antonella Succurro, Julie Maguire, Oliver Ebenhöh

**Affiliations:** Daithi O’Murchu Marine Research Station, Bantry, Co. Cork, Ireland; Institute for Quantitative and Theoretical Biology, Heinrich-Heine University, Düsseldorf, Germany; CEPLAS (Cluster of Excellence on Plant Sciences), Heinrich-Heine University, Düsseldorf, Germany

## Abstract

The pennate diatom *Phaeodactylum tricornutum* is a model organism able to synthesise industrially-relevant molecules. Large-scale monocultures are prone to bio-contamination, however, little is known about the identity of the invading organisms. To gain insight into the bacterial community associated with diatoms, we translated the complexity of a natural system into reproducible experiments where we investigated the microbiome of *P. tricornutum* cultures. The results revealed a dynamic bacterial community that changed over time and in differing media conditions. We propose a network of putative interactions between *P. tricornutum* and the main bacterial factions, which is translated into a set of ordinary differential equations constituting a computational dynamic model. The proposed mathematical model is able to capture the population dynamics, further supporting the hypothesised interactions. The interdisciplinary approach implemented provides a framework for understanding the dynamics of diatom-associated microbial communities, and provides a foundation for further systematic investigations of host-microbe interactions.

## Introduction

*Phaeodactylum tricornutum* is a diatom first described by Bohlin in 1897 when he found it in samples collected off the coast of Plymouth, United Kingdom. Diatoms belong to the Phylum *Heterokontophyta* and the Class *Bacillariophyceae* (Dangeard, 1933). They are the result of a secondary endosymbiotic event that took place around one billion years ago between a red alga (Rhodophyta) and a heterotrophic eukaryote (Bhattacharya *et al.*, 2007). Unlike most diatoms, which have the distinct ability to precipitate soluble silicic acid to form a silica cell wall, *P. tricornutum* has a poorly silicified cell wall and therefore does not have an obligate requirement for silicic acid (Montsant *et al.*, 2005; Martino *et al.*, 2007). *P. tricornutum* is found in coastal regions such as rock pools and estuaries where aquatic environmental parameters (salinity, temperature) vary greatly as a consequence of tidal changes and solar irradiation (Martino *et al.*, 2011). Its habitual characteristics, peculiar ability to form oval, fusiform, and triradiate cells, as well as its poorly silicified cell wall, have triggered a tremendous increase in scientific research on *P. tricornutum*. The genome sequencing of *P. tricornutum* was completed in 2008, and the subsequent generation of expressed sequence tag (ESTs) databases make *P. tricornutum* an excellent model organism (Montsant *et al.*, 2005; Martino *et al.*, 2007; Bowler *et al.*, 2008).

Driven by photosynthesis, *P. tricornutum* is able to synthesise a number of commercially relevant molecules, applicable to various industries. In aquaculture, *P. tricornutum* is used as feed for bivalve, echinoderm, crustacean and fish hatcheries (Ryther and Goldman, 1975; Tredici *et al.*, 2009). On average, 18% of the *P. tricornutum* biomass are lipids, making it a potential candidate for biofuel production (Kates and Volcani, 1966; Rebolloso-Fuentes *et al.*, 2001). Furthermore, *P. tricornutum* has the ability to produce the poly-unsaturated fatty acids (PUFA) eicosapentaenoic acid (EPA; 20:5n-3) and docosahexaenoic acid (DHA; 22:6n-3) in high proportions of the total fatty acid content (Siron *et al.*, 1989; Rebolloso-Fuentes *et al.*, 2001; Fajardo *et al.*, 2007). Marine-derived EPA and DHA, colloquially known as omega-3 PUFAs, are important in human nutrition with a vast number of health benefits (Yashodhara *et al.*, 2009). *P. tricornutum* is therefore an ideal source of omega-3 PUFAs for the pharma- and nutraceutical industries.

To fully exploit the industrial potential of *P. tricornutum* derived products, substantial amounts of microalgal biomass are required, preferably with low production costs. This is achieved by implementation of large-scale cultivation methods such as open raceway ponds and various types of photobioreactors. Microalgal cultivation methods rely on keeping monocultures of the desired species, especially if the final product is a bioactive molecule for human consumption (Mata *et al.*, 2010). Photobioreactors (PBRs) are closed systems that allow for the production of monoseptic cultures, fully isolated from potential contamination if cultivation protocols are followed correctly (Grima and Fernández, 1999). However, high operational costs of PBRs would increase production costs. The other option is open raceway ponds, which are simple open-air cultivation systems that have been in use since the 1950s (Chisti, 2007). They are highly susceptible to contamination, and unless the desired species is a halophile or thermophile (Parmar *et al.*, 2011), it is hard to maintain monocultures. Irrespective of the cultivation method, the establishment of unwanted organisms such as amoeba, ciliates, rotifers, bacteria, viruses, and other photosynthetic organisms in microalgal cultures, is a serious obstacle for large-scale microalgae cultivation (Day *et al.*, 2012; Wang *et al.*, 2013). Although much research is carried out in the field of microalgal culture upscaling, very little is known about the identity and characteristics of these invading organisms, responsible for microalgal culture ‘crashes’ which lead to loss of biomass, and therefore, loss of revenue.

The establishment of non-target organisms in microalgal cultures should not come as a surprise. Microalgae are not found in monoculture in nature and imposing such an environment is counterintuitive leading to unstable cultures. Rather than looking at these organisms as contaminants, understanding them could allow for the exploration of ‘synthetic ecology’ as a novel scaling up technique, a concept proposed by Kazamia *et al.*, 2012. The cornerstone of synthetic ecology is the Competitive Exclusion Principle, or Gause’s Law, which states ‘as a result of competition two species scarcely ever occupy similar niches, but displace each other in such a manner that each takes possession of certain peculiar kinds of food and modes of life in which it has an advantage over its competitor’ (Gause, 1934; Hardin, 1960). By ‘synthesising’ a community of organisms that fills every niche in the ecosystem (i.e. the microalgal culture) supporting the growth of the desired microalgae, we prevent the establishment of other, potentially harmful organisms in the culture, and optimise the utilisation of nutrients (Kazamia *et al.*, 2012).

In order for synthetic ecology to be a legitimate contender as a novel scaling up technique, greater understanding of species-specific interactions is required. Bacteria are present in all of the Earths’ biomes (Dykhuizen, 1998), and insight into the microorganisms (plankton) inhabiting our oceans was greatly improved by the three-year study abroad the schooner *Tara*. In May 2015, Sunagawa *et al.* published the metagenomics data from 243 samples collected from 68 unique locations during the *Tara* expedition. The data showed that 58.8% of the sequences belonged to bacteria, even though bacterial densities (10^5^ to 10^6^ per gram of seawater) in our oceans are orders of magnitudes less than those found in sediments (10^8^ cells per gram), humans (10^14^ cells per gram), or soil (10^9^ cells per gram) (Whitman *et al.*, 1998; Amin *et al.*, 2012). The data generated by the *Tara* project shows the sheer amplitude of genetic material belonging to bacteria, coupled with their co-existence with diatoms for more than 200 million years (Amin *et al.*, 2012), fuelled our interest in the microbiome of diatom cultures. Furthermore, in 1958, Provasoli suggested that bacteria can enhance the growth of algae (Provasoli, 1958). In the subsequent decades, species-specific studies have further corroborated his initial idea (Delucca and Mccracken, 1977; Suminto and Hirayama, 1997). Furthermore, Bruckner *et al.* showed an increase in growth of *P. tricornutum* when co-cultured with an Alphaproteobacterium strain as well as when cultured in the spent media of the bacteria (Bruckner *et al.*, 2011). A recent study conducted by Amin *et al.* shows a species-specific interaction between a coastal diatom, *Pseudo-nitzschia multiseries*, and a bacterial *Sulfitobacter* species (SA11), where the bacteria was shown to promote diatom cell division via secretion indole-3-acetic acid IAA, synthesised by the bacterium using diatom secreted and endogenous tryptophan. The IAA and tryptophan act as signalling molecules in this intricate diatom-bacteria relationship (Amin *et al.*, 2015).

With respect to the application in industry, the bacteria act as probiotics for the microalgae culture, just as bacterial probiotics have been successfully implemented in human diet by the pharma- and nutraceutical industries (Parvez and Malik, 2006), poultry industries (Kabir, 2009), and aquaculture industries (Qi *et al.*, 2009), to name a few. By identifying the bacterial community in non-axenic *P. tricornutum* cultures we can start to identify and characterise those that may have a beneficial role in the cultures. Subsequently, a suitable candidate to fill a certain niche in the hypothetical synthetic ecosystem could be chosen.

## Results

In order to translate the complexity of a natural system into a reproducible, systematic experimental approach, batch cultures of *Phaeodactylum tricornutum* (CCAP 1052/1B) were cultivated in two media conditions; (1) complete F/2 medium with the addition of sodium metasilicate as the source of silicon, and (2) minimal media with a source of nitrogen (NaNO_3_) and phosphorus (NaH_2_PO_4_.2H_2_O) at the same concentration as in the F/2 medium recipe. All *P. tricornutum* cultures were obtained from the Culture Collection of Algae and Protozoa (CCAP) based in Oban, Scotland (http://www.ccap.ac.uk/our-cultures.htm). All cultures are obtained non-axenic. Samples were taken at different stages of growth and subsequent barcoded 16S-V6-Next Generation Sequencing carried out. After the implementation of a stringent bioinformatics approach, the identity and abundance of the bacteria present in *P. tricornutum* cultures was revealed. The in the temporal evolution of the relative abundances of bacteria were used to infer a network of interactions between the diatom and the four dominant bacteria families, which was then translated into a mathematical model reproducing the community dynamics.

### Characteristics of *Phaeodactylum tricornutum* culture growth

The media composition was shown to have a significant effect on the growth characteristics of *P. tricornutum*. A significant difference (p=0.042, unpaired Wilcoxon signed rank) in the maximal cell density when *P. tricornutum* is cultivated in complete (9.3 x 10^6^ cells/mL) or minimal media (11.2 x 10^6^ cells/mL) was observed. The growth rates during the exponential phase in both cultures were µ_complete_ = 0.43 ± 0.07 d^−1^ and µ_minimal_ = 0.51 ± 0.04 d^−1^ respectively. In contrast, the death rates when the cultures ‘crash’ are δ_complete_ = 0.09 ± 0.02 d^−1^ and δ_minimal_ = 0.08 ± 0.04 d^−1^ respectively.

### Bacterial community profile of *Phaeodactylum tricornutum* cultures

In order to identify the bacteria present in the *P. tricornutum* cultures, the Ion Torrent™ barcoded Next Generation Sequencing protocol was used to sequence the bacterial gDNA (Quail *et al.*, 2012; Grada and Weinbrecht, 2013). The subsequent 16S rRNA gene sequences were clustered to defined Operational Taxonomic Units (OTUs) using a threshold of ≥97% sequence identity, most of which could be assigned to the genera level (Supplementary Figure S2). Of the 9727 OTUs identified, 8109 corresponded to known sequences in the SILVA database (v.118) (Quast *et al.*, 2013). The OTU abundance at the phylum level showed that 99.97% of all OTUs belonged to Proteobacteria, Bacteroidetes, Actinobacteria and Firmicutes (Figure 1). A comparison of the number of individual reads to the number of unique OTUs showed that the high number of reads per phyla is not the result of a single OTU (Supplementary Figure S3). OTUs with hits to known 16S *P. tricornutum* sequences were discarded.

**Figure 1.**
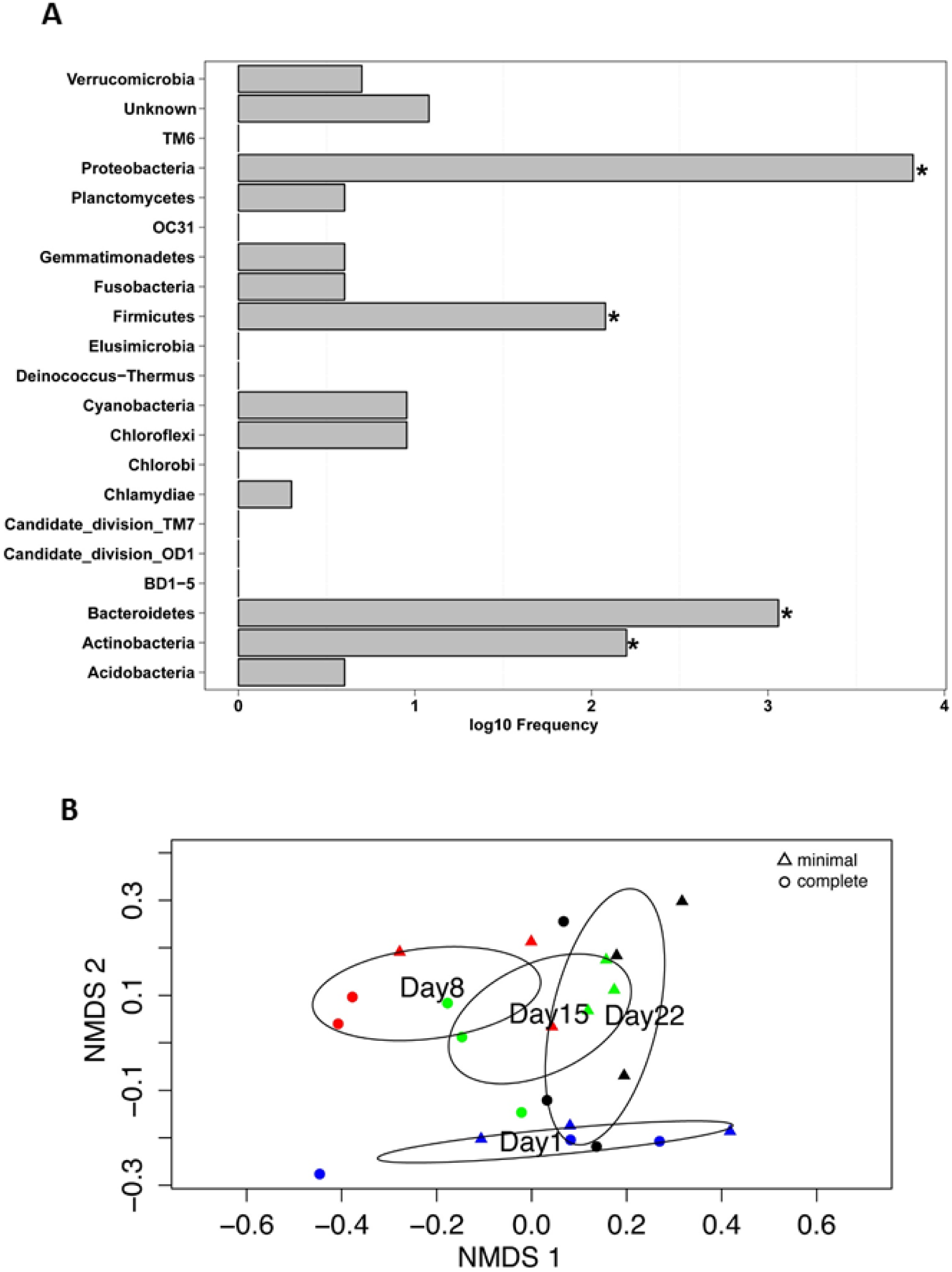
**(A) Distribution of Operational Taxonomic Unit (OTU) abundance (LOG scaled) within phyla from complete data set.** The bins marked with asterisks correspond to 99.97% of all which belong to Proteobacteria, Bacteriodetes, Actinobacteria and Firmicutes. **(B) Ordination plot of bacterial community in the two media conditions for all sampling points.** To compare the species composition between the different samples (days / media) we used a non-metric multidimensional scaling (NMDS) function based on generalised UniFrac distances (Chen *et al.*, 2012). Triangles and circles correspond to minimal media and complete media conditions, respectively. Blue represents Day 1. Red Day 8. Green Day 15. Black Day 22. The ellipses correspond to the 99% confidence interval to each group centroid.

Rarefaction curves were used to evaluate the alpha diversity in the different media conditions as well as at the different time points (Supplementary Figure S4). Species richness in both minimal and complete media was ~3000. Species richness over time remained between ~2400 and ~2600, with reduced species richness (~1300) on Day 8 (both minimal and complete media) possibly due to elevated levels of 16S *P. tricornutum* chloroplast reads which had to be omitted. Greatest species richness (~3000) was shown on Day 22. Overall, all datasets showed less increase in the number of unique species as the sample size increased, confirming adequate species richness in all culture conditions.

To compare the species composition between the different samples (days/media) we used a non-metric multidimensional scaling (NMDS) function based on generalised UniFrac distances (Chen *et al.*, 2012). We observed a clear divergence in the bacterial community in the two media conditions. Ordination based on the sampling day indicated that the bacterial community was dynamic with a clear divergence visible between Day 1 and the other three sampling days. Day 15 and 22 showed a slight overlap (Figure 2). An adapted version of PermanovaG was used to carry out permutational multivariate analysis of variance using multiple distance matrices which were previously calculated based on the generalised UniFrac distance (Chen *et al.*, 2012). The significance for the test was assessed by 5000 permutations. The results of the PermanovaG tests support the NMDS ordination, confirming a statistically significant effect in the bacterial community profile at the different sampling points and in the two media conditions whereas no significant effect was found in the experimental replicates (Supplementary Figure S5).

**Figure 2.**
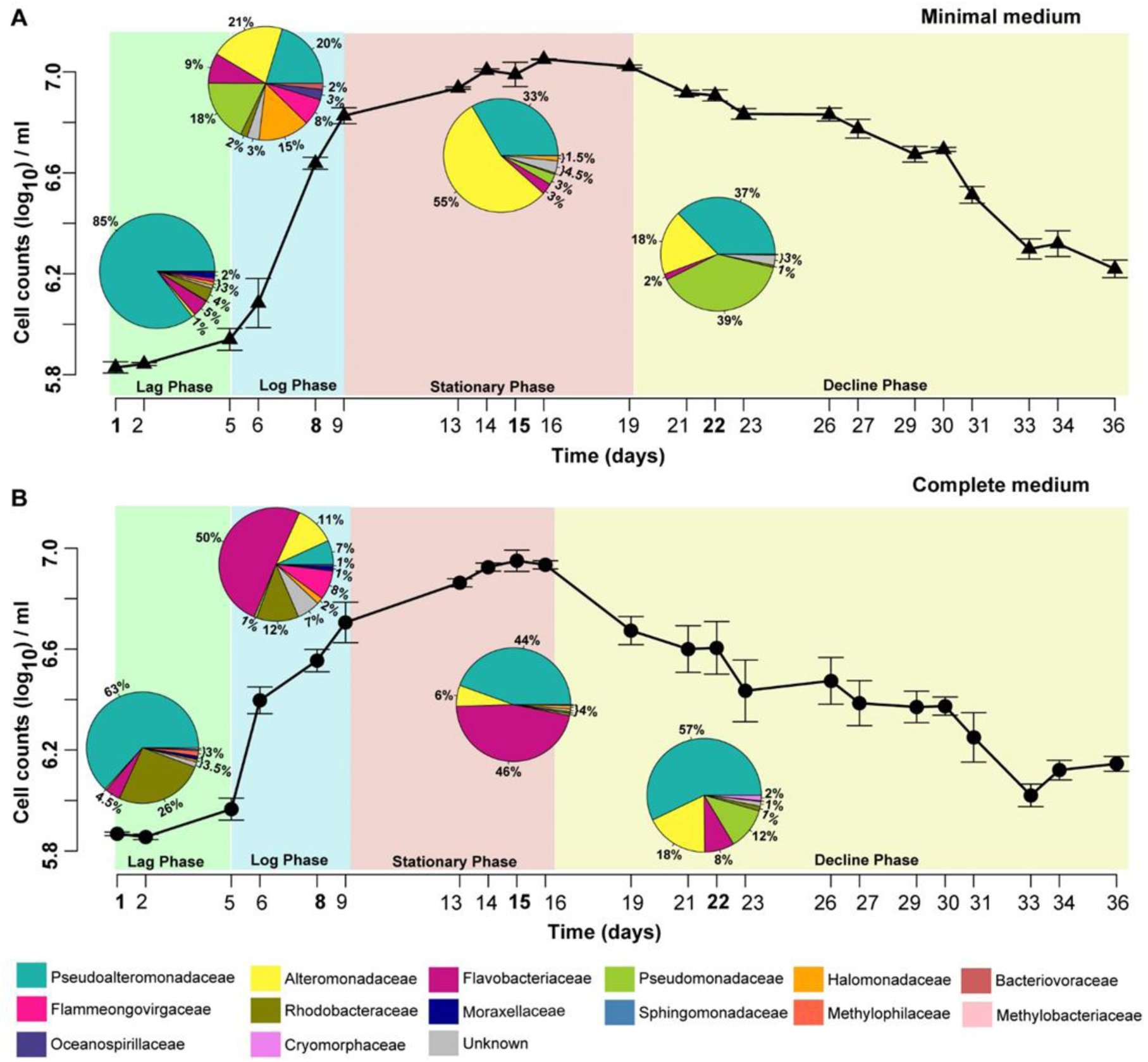
Bacterial community profile of *Phaeodactylum tricornutum* cultures over time and in differing media conditions. Both panels illustrate the growth of *P. tricornutum* (CCAP 1052/1B) over a 36 day period. The growth curves have been partitioned into lag (green), log (blue), stationary (red), and decline (yellow) phases. The abundance (%) of the ‘Top Ten’ bacterial families (corresponding colours described in the key) is depicted in pie charts on Days 1, 8, 15 and 22 in both media conditions. The existence of one dominant family at each investigated time point is a peculiar characteristic. In **minimal media (A)**, the lag phase of *P. tricornutum* growth is dominated by Pseudoalteromonadaceae (85%). However, during the log phase, a wide diversity of bacterial families is observed, with members of the Alteromonadaceae family (21%) beginning to dominate. During the stationary phase, a clear dominance of Alteromonadaceae species (55%) in the community can be observed. The decline phase, however, shows the Pseudomonadaceae (39%) as a dominant family, with Pseudoalteromonadaceae species (37%) increasing in abundance again. In **complete media (B)**, the lag phase is also dominated by Pseudoalteromonadaceae (63%). During the log phase, 50% of the community is composed of members of the Flavobacteriaceae family, with the other 50% distributed amongst a number of different families. Flavobacteriaceae (46%) remain high in abundance during the stationary phase, with Pseudoalteromonadaceae species (44%) beginning to increase in abundance again. As for minimal media (A), Pseudoalteromonadaceae (57%) show clear dominance of the community during the decline phase.

### Effect of temporal evolution and media composition on the bacterial community profile

We compared the bacterial community profiles over time and in the different media conditions at the family level so as to avoid diluting the signal of the less abundant genera. Supplementary Figure S6 shows no dynamical difference within the genera that cannot be observed at the family level. By investigating the bacterial community dynamics at the family level, we also include taxonomical information that is unavailable at the genus level.

Overall, the families over-represented in all samples are Pseudoalteromonadaceae, Alteromonadaceae, Flavobacteriaceae and Pseudomonadaceae. Figure 2 illustrates the temporal evolution of the bacterial community in both minimal and complete media with a unique composition at each time point. A remarkable feature is that at all investigated time points there exist one or two dominant families.

### Bacterial community in complete media

Members of the Pseudoalteromonadaceae family were highly abundant when *P. tricornutum* cell densities are low (63% and 57% on Day 1 and Day 22, respectively). Flavobacteriaceae species dominated (50%) when the *P. tricornutum* culture is growing exponentially (Day 8). Day 15, when *P. tricornutum* cell densities are at their highest, shows co-dominance of both Flavobacteriaceae (46%) and Pseudoalteromonadaceae (44%).

### Bacterial community in minimal media

Similarly, in the minimal media, members of the Pseudoalteromonadaceae family were highly abundant when *P. tricornutum* cell densities are low. However, on Day 22 Pseudomonadaceae (39%) and Pseudoalteromonadaceae (37%) are both overrepresented. When the *P. tricornutum* culture is growing exponentially (Day 8) a cluster of Families dominate; namely Alteromonadaceae (21%), Pseudoalteromonadaceae (20%), Pseudomonadaceae (18%), Halomonadaceae (15%) and Flavobacteriaceae (9%). When the cell density of *P. tricornutum* peaks (Day 15), the Alteromonadaceae species take over (55%).

The bacterial communities within the two media conditions on Day 1 are more closely related than the communities on days 8 and 15 (see Table S2 for generalised UniFrac distances). As the cultures begin to ‘crash’ (Day 22), the bacterial communities in the two media conditions increase in similarity again.

In general, the main families identified show a distinct pattern of disappearance and regeneration within the bacterial community. In the complete media, Pseudoalteromonadaceae species start at 63% (Day 1), drops in abundance to 7% (Day 8) then recovers to 57% (Day 22). Flavobacteriaceae species, in complete media, start at 4.5% (Day 1), increases in abundance to 50% (Day 8), and then falls back to 8% (Day 22). In the minimal media, Alteromonadaceae species have an abundance of only 1% (Day 1), peaks at 55% (Day 15), and decreases down to 18% (Day 22).

### Mathematical model

The dynamic changes of the bacterial communities associated with *P. tricornutum* at different growth stages led us to the formulation of a network of bacteria - diatom interactions. In order to test its plausibility, we developed a qualitative mathematical model starting from few key assumptions about nutrients availability and metabolite exchange between the organisms involved, i.e. *P. tricornutum* and general representatives of the four most abundant bacteria families Pseudoalteromonadaceae, Alteromonadaceae, Flavobacteriaceae and Pseudomonadaceae.

In Figure 3A and B we show the simulation results from the model performed in complete media and minimal media conditions, respectively, with experimental data superimposed. The top panel shows biomasses of the five organisms (data available only for the diatom), the bottom panel shows relative bacteria abundance versus time (biomass divided by total bacteria biomass). The figures show that the model is able to reproduce the main features of the bacterial community dynamics, like the disappearance and return of Pseudoalteromonadaceae in complete media and the peak of Alteromonadaceae at the end of the diatom’s exponential growth phase in minimal media. Because of the qualitative nature of the model, units are arbitrary and the parameters used for simulation do not claim any biological significance (see Supplementary Model Information for more details).

**Figure 3A and B.**
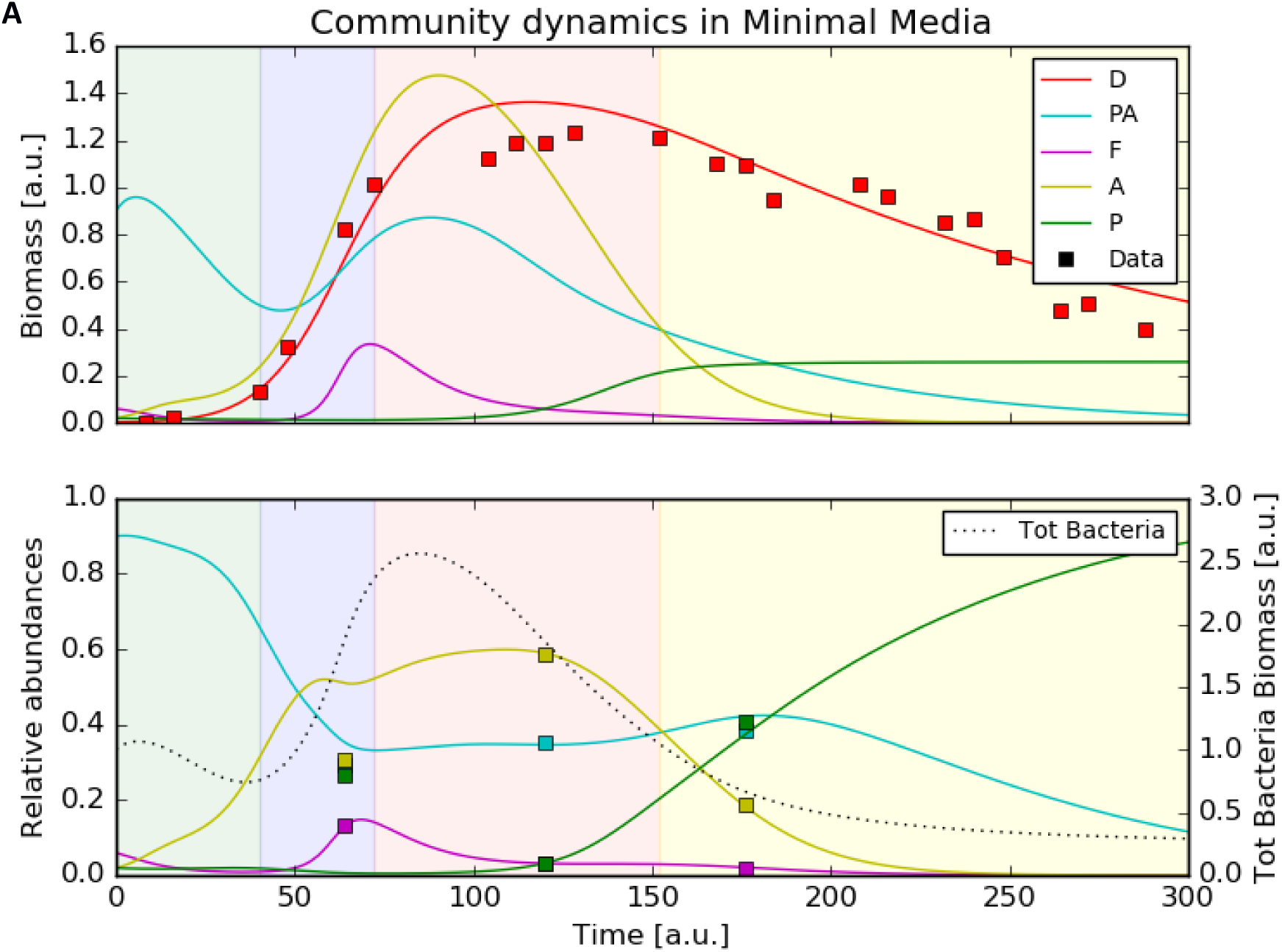
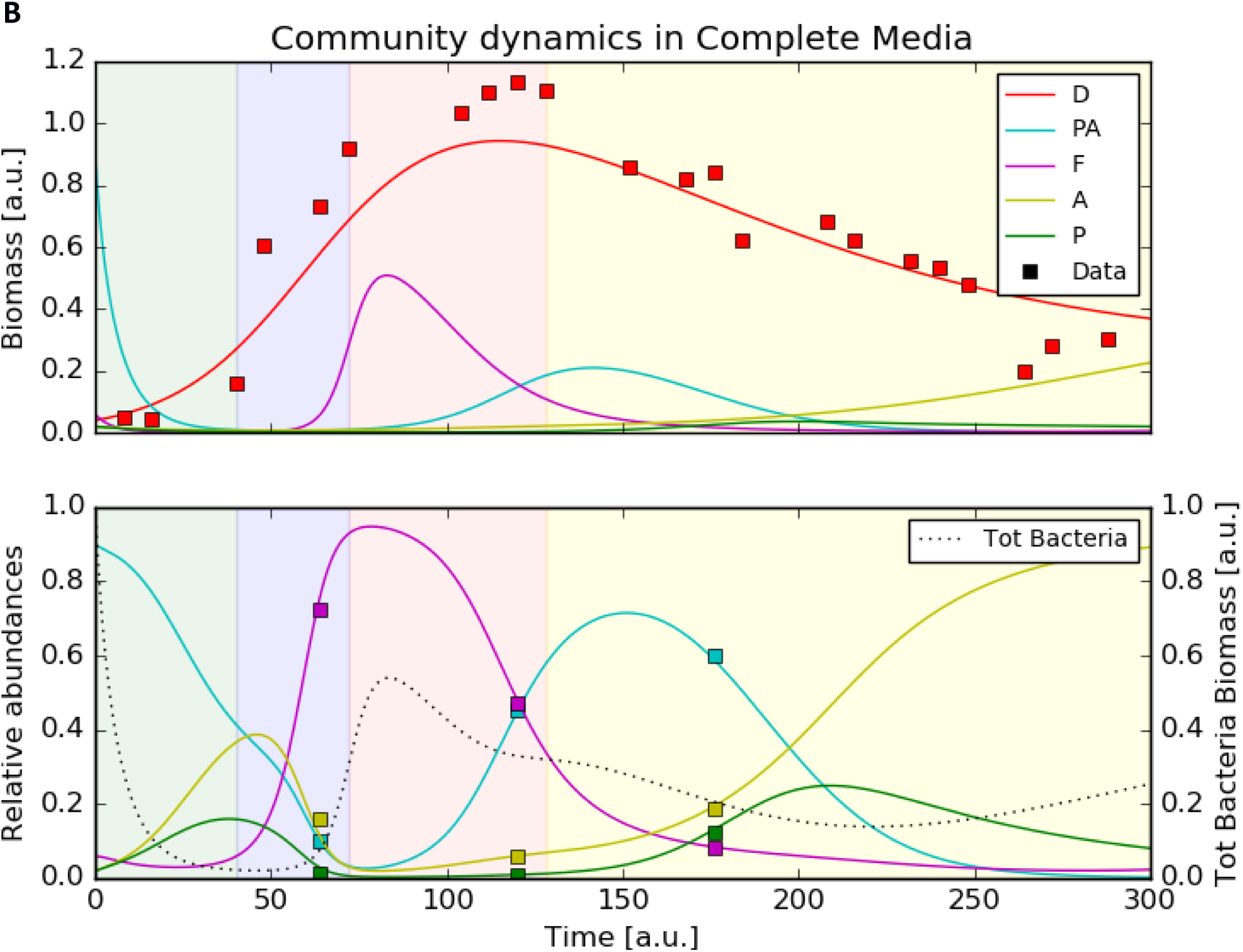
Simulation results (lines) and experimental data (squares) for communities of *P. tricornutum* (D), Pseudoalteromonadaceae (PA), Flavobacteriaceae (F), Alteromonadaceae (A) and Pseudomonadaceae (P) in minimal (A) and complete (B) media conditions. The top panel shows the biomass time course (arbitrary units) for the five organisms and the rescaled data points (squares) for the *P. tricornutum*. The bottom panel shows the variations in relative abundances of the four bacteria (single bacteria biomass/total bacteria biomass) over time and the three sets of data points from the sequencing analysis (the first data point is used as starting condition at time 0). Also shown in the bottom plot (dotted line, right y-axis) is the total bacterial biomass in arbitrary units.

## Discussion

In nature, *Phaeodactylum tricornutum* is not an isolated sovereign entity impassive to its environment including other organisms. In fact, it is part of a complex ecosystem, which is poorly understood. To reduce the complexity of a natural system, but nevertheless to gain valuable insights into the dynamics of the bacterial communities associated with diatoms, we investigated here non-axenic cultures of laboratory strains of *P. tricornutum*. Our results showed the trends of the bacterial community dynamics during the batch growth of the *P. tricornutum*.

To progress towards the goal of creating a synthetic community, an in-depth understanding of the naturally occurring diatom-bacterial interactions, which are predominantly based on a ‘biological barter trade system’ between diatoms and bacteria – where substances such as trace metals, vitamins, and nutrients (nitrate, phosphate, silicate, carbon) are traded – is necessary. Based on our findings and additional insights from previous studies on diatom-bacterial interactions as well as on existing characterisation of known species from each family, we will postulate the role of the particular bacterial families in the *P. tricornutum* cultures. From this we will derive a mathematical model with the aim of reproducing the dynamical evolution of the community composition over time.

The growth dynamics of *P. tricornutum* in the two media conditions showed an accelerated ‘culture crash’ in the complete media compared to the minimal media, which suggests a more stable culture in the minimal media (Figure 2). Simultaneously, the dynamics of the bacterial community reveals that the community in the minimal media increases in complexity over time. The link between ecosystem complexity and stability based on theoretical and experimental data has been debated by ecologists for over half a century (MacArthur, 1955; Elton, 1958; Gardner and Ashby, 1970; Pimm, 1984). Our observations are in agreement with more recent hypotheses indicating that diversity generally increases the stability of an ecosystem (McCann, 2000).

### Prospective role of central bacterial families

The putative roles of each of the dominant families are illustrated in Figure 4. The presence of **Pseudoalteromonadaceae** species is not unexpected as members of this family have been isolated from coastal, open and deep-sea waters, sediments, marine invertebrates, as well as marine fish and algae (Ivanova *et al.*, 2004). The Pseudoalteromonadaceae family has three genera, namely *Pseudoalteromonas*, *Algicola* and *Psychrosphaera* (Rosenberg *et al.*, 2014, 28). Several species of Pseudoalteromonadaceae are reported to possess antibiotic properties with bactericidal effects (Bowman, 2007). For example, concentrated supernatant of a marine bacterium *Pseudoalteromonas sp.* strain A28 contained various enzymes including proteases, DNases, cellulases, and amylases, capable of causing the lysis of the diatom *Skeletonema costatum* (Lee *et al.*, 2000). Species of Pseudoalteromonadaceae are also capable of producing cold-adapted enzymes (Venkateswaran and Dohmoto, 2000; Chen *et al.*, 2007; Khudary *et al.*, 2010; Lu *et al.*, 2010; Albino *et al.*, 2012; He *et al.*, 2012). Pseudoalteromonadaceae species can produce extracellular polymeric substances allowing them to colonise surfaces, enhancing nutrient uptake whilst limiting diffusion of particular substances across the cell membrane (Holmström and Kjelleberg, 1999). The ability of Pseudoalteromonadaceae species to suppress the growth of competing bacteria could explain the dominance of Pseudoalteromonadaceae in almost all cultures irrespective of media composition, particularly when *P. tricornutum* abundance is limited (Figure 2, Days 1 and 22). *P. tricornutum* on the other hand, may protect other bacterial community members from the bacteriolytic ability of Pseudoalteromonadaceae by producing specific antibacterial compounds themselves. Desbois *et al.* showed that *P. tricornutum* excreted bacteriolytic fatty acids such as eicosapentaenoic acid (EPA; 20:5n-3), nucleotides, peptides, and pigment derivatives that can eliminate unwanted competition for nutrients such as organic phosphates from certain bacteria (Desbois *et al.*, 2009).

**Figure 4.**
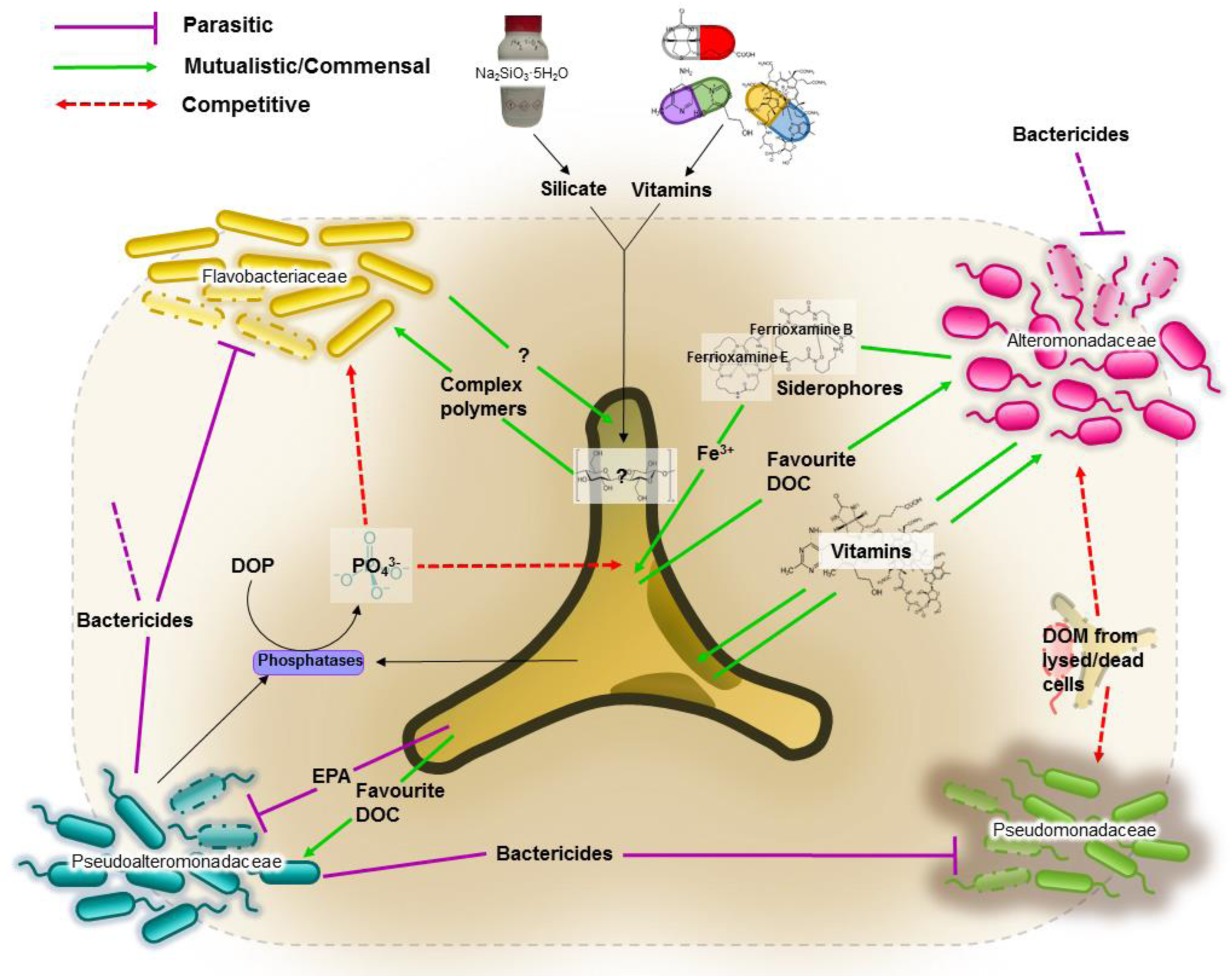
Network of putative interactions between *Phaeodactylum tricornutum* and identified bacterial families. The dotted grey line depicts the **‘phycosphere’**; a term coined by Bell and Mitchell in 1972 as an aquatic equivalent of the ‘rhizosphere’, denoting the region extending outwards from the algal cell in which bacterial growth is stimulated by extracellular products of the alga (Bell and Mitchell, 1972). **Bactericidal Effects.** Several species of the Pseudoalteromonadaceae family have been reported to possess bactericidal effects (Bowman, 2007). *P. tricornutum*, however, can excrete fatty acids (such as eicosapentaenoic acid or EPA), nucleotides, peptides, and pigment derivatives to protect themselves against opportunistic attack or pathogenic damage (Desbois *et al.*, 2009). **Iron.** Siderophores are a group of iron scavengers that act by chelating iron (III). Siderophores are produced and excreted by bacteria, and some cyanobacteria, which then reuptake the siderophores with bound iron (III) via outer-membrane transporters that are siderophore-specific (Vraspir and Butler, 2009). Diatoms are not known to produce siderophores (Soria-Dengg and Horstmann, 1995; Amin *et al.*, 2009). However, based on genome sequence analyses, the presence of a gene orthologue of a bacterial ferrichrome binding protein suggests the possibility of iron (III)-siderophore utilisation by *P. tricornutum*. Furthermore, it was shown that *P. tricornutum* was able to uptake siderophores ferrioxamines B and E (Soria-Dengg and Horstmann, 1995). **Vitamins.** Prokaryotes are thought to be the main producers of B vitamins (Provasoli, 1963; Provasoli and Carlucci, 1974). Although *P. tricornutum* does not require cobalamin, thiamine and biotin (Croft *et al.*, 2006), production of organic compounds such as EPA can by considerably enhanced by the bioavailability of co-factors such as cobalamin (Yongmanitchai and Ward, 1991). This provides the basis for potential mutualistic interactions. For example, Alteromonadales, dominant in our cultures, are thought to be capable of producing B vitamins (Sañudo-Wilhelmy *et al.*, 2014). **Dissolved Organic Carbon (DOC).** It is estimated that up to 50% of carbon fixed via phytoplankton-mediated photosynthesis is utilised by marine bacteria (Azam *et al.*, 1983), mainly as DOC compounds, defined as the organic material <0.7µm in size (Stocker, 2012). DOC from diatoms originates either from live cells or recently lysed or grazed cells, which determines the type of DOCs available, and therefore likely determining the bacterial consortia associated with the diatom (Amin *et al.*, 2012). **Dissolved Organic Phosphate (DOP).** Both diatoms and bacteria primarily utilise orthophosphate as a source of phosphorus. However, to access phosphate from DOP compounds, both diatoms and bacteria developed mechanisms such as the excretion of enzymes, including phosphatases, to release orthophosphate (PO_4_^3-^) from DOP. The mechanism is not species-specific, which consequently means the ‘free’ orthophosphates can be acquired by any organism (Persson *et al.*, 1988).

The **Alteromonadaceae** family consists of 16 (yet annotated) named genera (LPSN, 2016a) found predominantly in marine environments (Rosenberg *et al.*, 2014, 5). Members of this family were isolated from nutrient-rich environments such as coastal, open, and deep-sea waters, sediments, marine invertebrates and vertebrates, algae, and temperate and Antarctic marine environments (Ivanova and Mikhaĭlov, 2001). They are able to utilise a vast array of compounds as carbon sources; from glucose to glycerol (Rosenberg *et al.*, 2014, 5). Members of this family are known siderophore producers (Reid and Butler, 1991; Holt *et al.*, 2005; Amin *et al.*, 2009). Greek for ‘iron carrier’, siderophores are a group of iron scavengers that act by chelating iron (III) that are produced and excreted by bacteria, and some cyanobacteria, which then reuptake the siderophores with bound iron (III) via outer-membrane transporters that are siderophore-specific (Vraspir and Butler, 2009). Iron acquisition is essential for biological processes such as photosynthesis, respiration and nitrogen fixation. Most bioactive trace metals, including iron, exist at nanomolar (10^−9^ M) to picomolar (10^−12^ M) concentrations in our oceans, approximately one-millionth of the intracellular concentration in diatoms (Bruland *et al.*, 1991; Morel and Price, 2003). Diatoms are not known to produce siderophores (Soria-Dengg and Horstmann, 1995; Amin *et al.*, 2009) but previous studies have shown that diatoms can use siderophores as an iron source (Soria-Dengg *et al.*, 2001). However, based on genome sequence analyses, the presence of a gene orthologue of a bacterial ferrichrome-binding protein suggests the possibility of iron (III)-siderophore utilisation by *P. tricornutum* (Soria-Dengg and Horstmann, 1995). No trace metals, including iron (III), were provided to minimal media cultures. However, natural seawater may contain minute traces of bioactive trace metals. The high abundance of Alteromonadaceae in the minimal media suggests a potential supportive role in sequestering traces of iron (III) that may be present in the sterile natural seawater to the *P. tricornutum* (Figure 2). This is further supported by the very low level of Alteromonadaceae in the complete media (11% in complete media compared to 55% in minimal media, both on Day 15) where the culture has been supplied with 11.7 µM of iron (III) chloride hexahydrate.

**Flavobacteriaceae** are members of the Bacteroidetes phylum and include over 120 genera (LPSN, 2016b) found in soil, sediments and seawater (see (Yoon *et al.*, 2015) for further references). Flavobacteriaceae belong within the Cytophaga-Flavobacterium cluster which has been shown to account for more than 10% of the total bacterial community in coastal and offshore waters (Glöckner *et al.*, 1999; Abell and Bowman, 2005; DeLong *et al.*, 2006). Members of Flavobacteriaceae are proficient degraders of various biopolymers such as cellulose, chitin and pectin (Manz *et al.*, 1996; Kirchman, 2002). They were shown to be omnipresent during phytoplankton blooms, and their preference for consuming more complex polymers rather than monomers suggests an active role in the processing of organic matter during these blooms (Cottrell and Kirchman, 2000; Pinhassi *et al.*, 2004). Although the exact mechanisms behind them are not perfectly understood, algal blooms are a consequence of exponential growth of phytoplankton (Smayda, 1997). In this respect, the phase of exponential growth of *P. tricornutum* in complete media, when our results showed highest abundance of Flavobacteriaceae, is the artificial equivalent of an algal bloom of *P. tricornutum* (Figure 2). In the minimal media, the abundance of Flavobacteriaceae remains very low; at its maximum on Day 8 it only accounts for 9% of the total bacterial community. Members of the Flavobacteriaceae family could be more demanding than other bacteria that require lower nutrient levels to thrive. It is estimated that up to 50% of carbon fixed via phytoplankton-mediated photosynthesis is utilised by marine bacteria (Azam *et al.*, 1983), mainly as Dissolved Organic Carbon (DOC) compounds, defined as the organic material <0.7µm in size (Stocker, 2012). DOC from diatoms originates either from live cells or recently lysed or grazed cells, which determine the type of DOCs available, and therefore are likely to influence the bacterial consortia associated with the diatom (Amin *et al.*, 2012). This suggests a dynamic complexity within the bacterial consortia based solely on the type of DOC available. Members of the Flavobacteriaceae family might possess the genetic ability to utilise specific DOC produced by *P. tricornutum* grown in complete media.

**Pseudomonadaceae** are an extraordinarily diverse family of bacteria found in almost all habitats on Earth; in soils, freshwater as well as marine environments, as well as plant and animal-associated pathogens (Starr *et al.*, 1981, 58). Species from the *Pseudomonas* genus are the best studied of the Pseudomonadaceae family, whose sheer genetic diversity explains the ability to thrive in such a wide range of environments (Anzai *et al.*, 2000). Marine isolates from the *Pseudomonas* genus have been shown to produce a wide range of bioactive compounds, many of which exhibit antibacterial as well as antiviral properties (see (Isnansetyo and Kamei, 2009) for further references). Our results, indeed show an elevated level of Pseudomonadaceae OTUs evident on Day 22 of the complete media cultures, and on Days 8 and 22 of the minimal media cultures. The increased presence of Pseudomonadaceae when *the P. tricornutum* culture has ‘crashed’ could be attributed to its ability to produce antibacterial compounds allowing members of this family to begin to thrive in the community through inhibition of its competitors. Given its exceptional genetic diversity, and thus, its metabolic versatility, allows for members of Pseudomonadaceae to be truly saprophytic; providing a hypothetical explanation of its abundance we could measure when the *P. tricornutum* cultures crash (Figure 2, Day 22 in both media conditions).

### Mathematical Model

We observed that the bacterial community associated with *Phaeodactylum tricornutum* cultures changed over time, correlating with the growth and subsequent crashing of the diatom cultures. The bioavailability or absence of vitamins, trace metals and silicon, as well as nutrients or bactericidal substances can alter the bacterial community. We built a mathematical model based on simple assumptions extracted from the putative roles we assigned to the dominant bacterial families (see Figure 4) and applied them to standard methods for modelling population dynamics. In particular, we introduced growth limitation from nutrients/micronutrients, as well as from bactericidal-induced death. An ordinary differential equation model cannot, of course, capture mechanisms such as metabolic shifts caused by changes in the environment such as the supplementation of minimal or complete media. Therefore, we did not implement a unique set of parameters for the model in the two conditions. The current qualitative model provides an important proof-of-concept to emphasise the validity of our assumptions, and serves as the motivation for further research bringing the model to a quantitative, predictive level. Indeed, mathematical models are powerful tools towards the goal of synthetic community establishment and control, and the model parameters can be experimentally measured to bring predictive power to the simulations.

### Concluding remarks

We postulate that a role within the community can be filled, not by one specific species of bacteria, but rather a number of bacterial species capable of carrying out said role. Which bacteria fill the role is dependent upon the environmental characteristics and the prevailing needs of the community as a whole at any given time. If a niche is unfilled, bacteria with the ideal metabolic functionality will seize the opportunity and thrive within that niche. The absence of certain micronutrients creates a new niche that can be filled by a certain unique bacterial faction.

Further work is necessary to explore the hypotheses postulated in the Discussion section. This can be achieved by carrying out systematic co-culture experiments with culturable members of the bacterial families of interest. The role of each representative of the bacterial families can be identified by carrying out subsequent –omics studies, which provide a holistic view of the genes (genomics), mRNA (transcriptomics), proteins (proteomics) and metabolites (metabolomics) in a specific biological sample in a non-targeted and non-biased manner (Horgan and Kenny, 2011). The resulting experimental measurements will allow the dynamic model presented here to develop from qualitative to quantitative, providing a powerful predictive tool for culture monitoring such as predicting harvesting point based on the bacterial community.

## Materials and methods

### Strains and culture conditions

All *Phaeodactylum tricornutum* cultures were obtained from the Culture Collection of Algae and Protozoa (CCAP) based in Oban, Scotland (http://www.ccap.ac.uk/our-cultures.htm). All cultures are obtained non-axenic. Based on previous experimental evidence (unpublished data), the *P. tricornutum* strain CCAP1052/1B displayed optimal growth in 5L cultures. *P. tricornutum* was cultured in Guillard’s medium for diatoms (F/2 + Si) in filtered natural seawater chemically sterilised using sodium hypochlorite and sodium thiosulphate pentahydrate. *P. tricornutum* was grown in two media conditions; (1) complete F/2 medium with the addition of sodium metasilicate as the source of silicon, as per Guillard and Ryther, 1962; Guillard, 1975, and (2) minimal media with a source of nitrogen (NaNO_3_) and phosphorus (NaH_2_PO_4_.2H_2_O) at the same concentration as in the F/2 medium recipe. Recipe was obtained from the Culture Collection of Algae and Protozoa website (see http://www.ccap.ac.uk/pdfrecipes.htm). All cultures were grown in hanging 5L polyethylene bags with a ‘V’ shaped bottom prepared using a heat sealer (Supplementary Figure S1). All cultures had a modified aeration system provided by a 10ml pipette attached to the main pressurised air supply via 0.2 µm sterile air filters. A modified access port was created to allow for sampling and measurement of environmental parameters. Cultures were kept at 18-20°C and 24hr light at an average of 132.3 µmol m^−2^ s^−1^ using Phillips TL-D 58W 33-640 M9 fluorescent tube lights. All cultures, irrespective of media condition, were inoculated with 250ml from the same 5L stock culture of actively growing non-axenic *P. tricornutum*.

### Growth measurements

Growth was monitored every 24 to 48 h using a light microscope and carrying out cell counts of each culture in quadruplicates for each culture. During the cell counts the ratios of the four different morphotypes (oval, fusiform, triradiate and cruciform) were recorded, and descriptions of each culture noted. Samples of each culture were subsequently taken using a sterile 10ml syringe and placed in 50ml Falcon centrifuge tubes and placed in −20°C freezer.

### Genomic DNA extraction

All samples from Day 1, 8, 15, and 22 were thawed in a water bath set at 25°C. As per de Gouvion Saint Cyr *et al.*, 2014, samples were centrifuged for 5mins at 2000g to gather the *P. tricornutum* in the pellet while particles such as debris, other organisms, bacteria, and soluble substances remain in the supernatant. Because the bacteria might be attached to the *P. tricornutum* cells in the pellet, the pellet was washed with deionised water and then centrifuged for 5mins at 2000g. This was repeated twice. Genomic DNA extraction was carried out in the Aquaculture and Fisheries Development Centre and University College Cork. The Mo Bio’s PowerWater^®^ DNA Isolation Kit (catalogue no. 14900-100-NF) was utilised to carry out the genomic DNA extraction. The protocol provided with the kit was followed. Presence of gDNA was detected by running a 1% agarose-ethidium bromide gel with 72 wells. The samples were sent on dry ice to Heinrich Heine University, Düsseldorf, for the V6 16S sequencing.

### Barcoded 16S-V6-Next Generation Sequencing

Ion Torrent™ barcoded Next Generation Sequencing protocol was used to sequence the bacterial gDNA (Quail *et al.*, 2012; Grada and Weinbrecht, 2013). Amplification of the V6 hyper variable region of 16S rRNA with forward and reverse primers (Supplementary Table S2) was carried out. Ion Reporter™ software assembles all the raw sequencing data and sorts all the reads using the unique sample-specific barcode sequences and removes them from the reads. The outcome is raw FASTQ files which are ready for analysis using bioinformatics tools.

### Bioinformatics

A total of 87,077,374 reads were identified. The smallest sample was just over 1 million reads; the largest sample was just under 10 million reads. The sequencing data was subjected to a pipeline adapted and modified from Pylro *et al.*, 2014. Primers were trimmed with fastq-mcf (version 1.04.807) (Aronesty, 2011), the resulting sequences were quality filterted and clustered into OTUs (operational taxonomic units) with usearch (version 8.0.1517; 32Bit – opensource) (Edgar, 2010, 2013). Taxonomy assignment was done by QIIME (version 1.9.0) (Caporaso *et al.*, 2010) with the implemented uclust classifier based on 97% sequence identity to the reference 16S sequences from SILVA 111 database (Quast *et al.*, 2013). Statistical analyses were performed in R (R Development Core Team, 2015). The complete protocol containing all processing steps is available on https://github.com/QTB-HHU.

### Modelling approach

Population dynamics models have been developed since quite some time (Verhulst, 1838; Lotka, 1925; Volterra, 1926) spanning the broad fields of ecology, epidemiology and economics. Starting from our understanding of the organism-to-organism interactions, we developed a dynamic model consisting of 13 ordinary differential equations and including 56 (55 free) parameters. The parameters are fitted using a genetic algorithm (Mitchell, 1996) which is run in different steps to optimise the fit of *P. tricornutum* growth and/or the bacteria relative abundances to the experimental data in evolving system conditions (see Supplementary Model Information). The model is written in Python (Python Software Foundation, https://www.python.org/) and is available on GitHub (https://github.com/QTB-HHU/communityODE) with instructions and scripts for running.

## Acknowledgements

This research was funded by the Marie Curie Initial Training Network project ‘AccliPhot’ (grant agreement number PITN-GA-2012-316427). Genomic DNA extraction was carried out at the Aquaculture and Fisheries Development Centre, University College Cork, Ireland (funded by Beaufort Marine Research Award in Fish Population Genetics funded by the Irish Government under the Sea Change Programme). Barcoded 16S-V6-Next Generation Sequencing was carried out by the Genomics and Transcriptomics Laboratory at Heinrich-Heine University, Düsseldorf, Germany. OP and OE are funded by Deutsche Forschungsgemeinschaft, Cluster of Excellence on Plant Sciences, CEPLAS (EXC 1028).

## Supplementary Material

**Figure S1.**
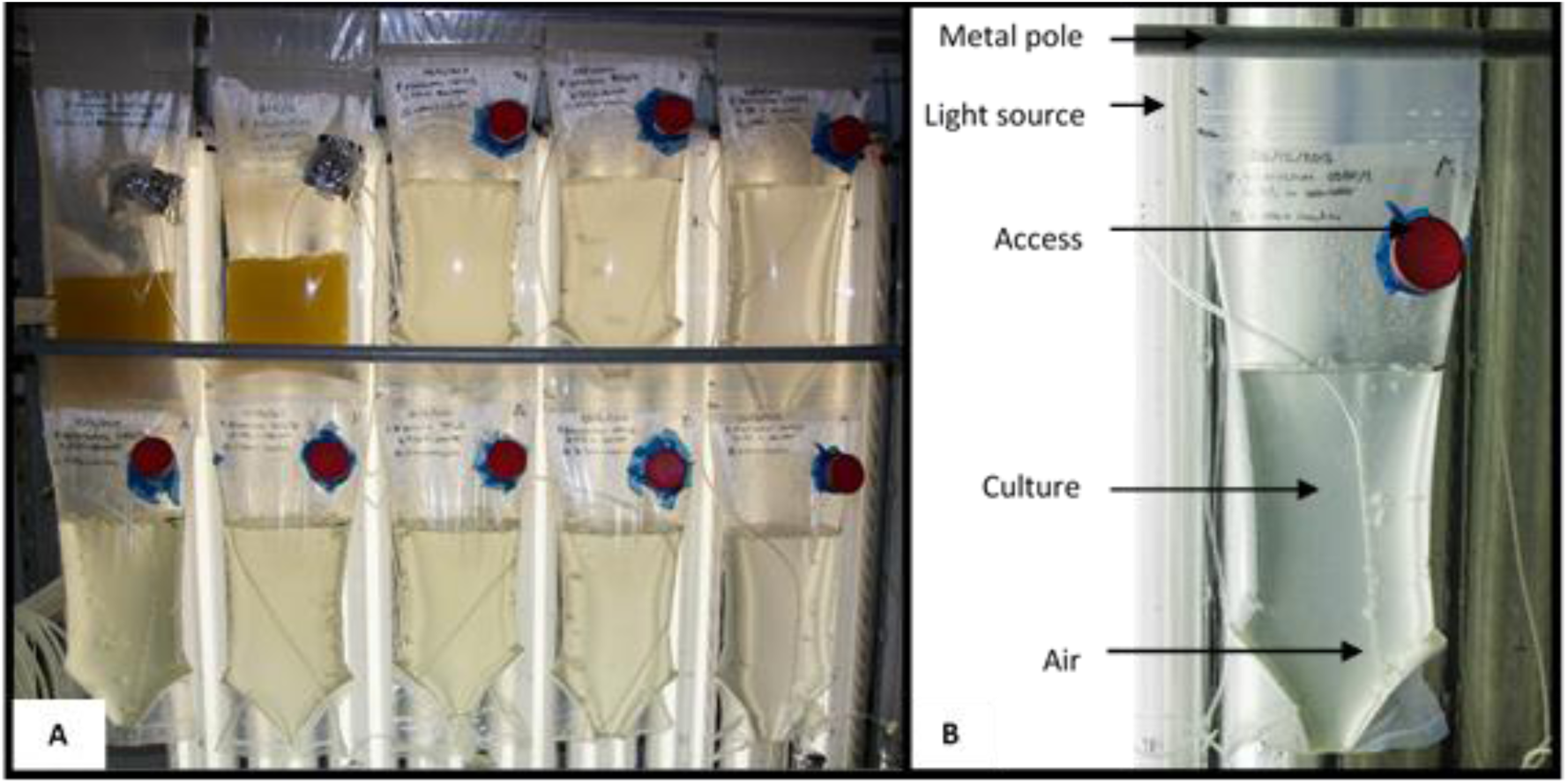
Non-axenic *Phaeodactylum tricornutum* culture set up. 5L polyethylene bags with a ‘V’ shaped bottom were created using the heat sealer machine. The bags were then rinsed and filled with 5L of filtered seawater. Afterwards each bag was sealed and hung approximately 30 cm from the light source. A small incision was made to insert the aeration tubing. This consists of a 10ml pipette attached to silicon tubing which is attached to a sterile air filter connecting it to the main air supply. A modified access port was created to take samples and measure the environmental parameters (Photographs courtesy of Maria Rubio Bernal)

**Figure S2.**
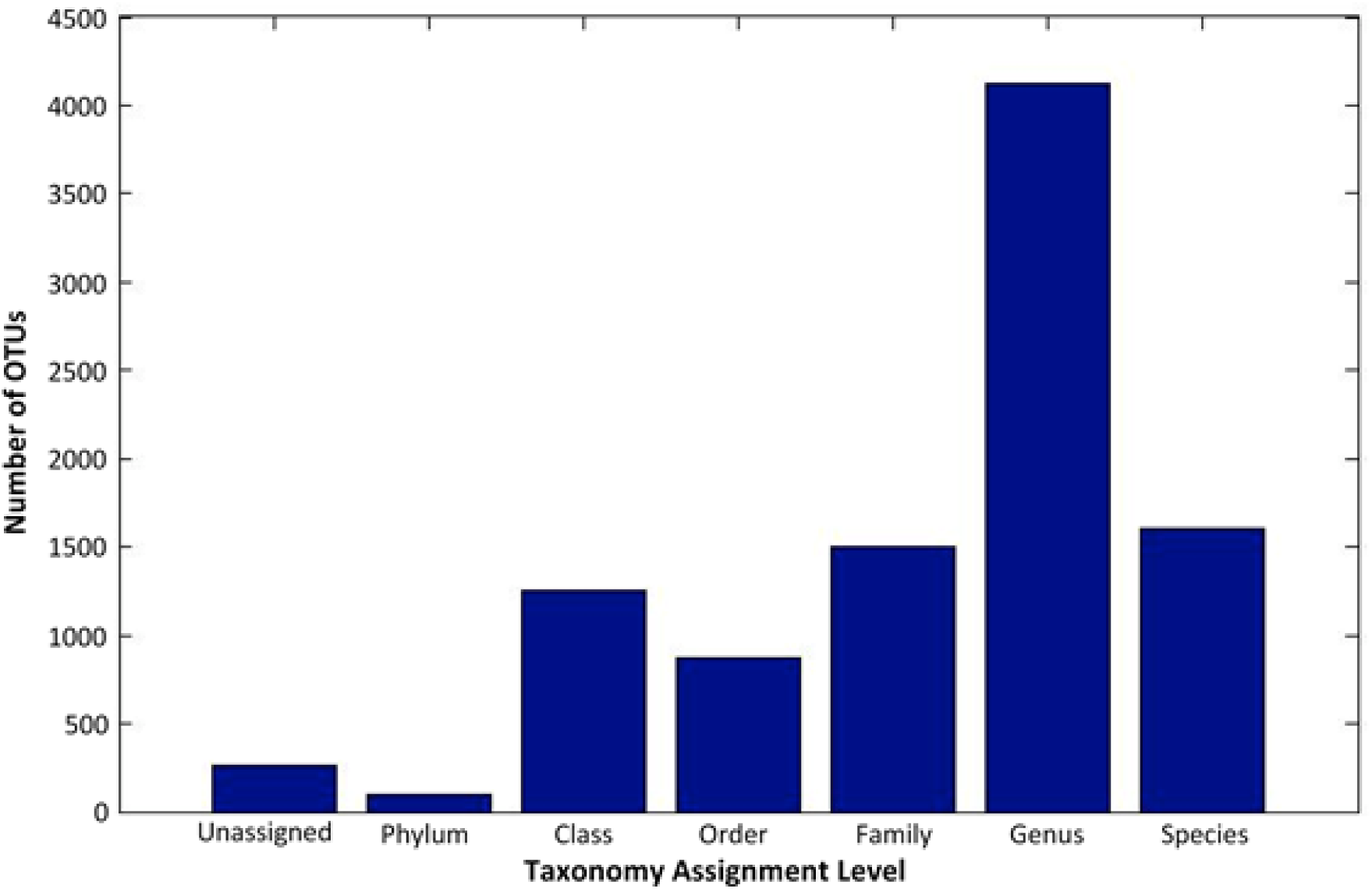
Operational Taxonomic Unit (OTU) Taxonomy Assignment Level. The 16S rRNA gene sequences were clustered to defined Operational Taxonomic Units (OTUs) at ≥97% sequence identity. Most OTUs could be assigned to the genera level, using the SILVA database (v.118) (Quast *et al.*, 2013).

**Figure S3.**
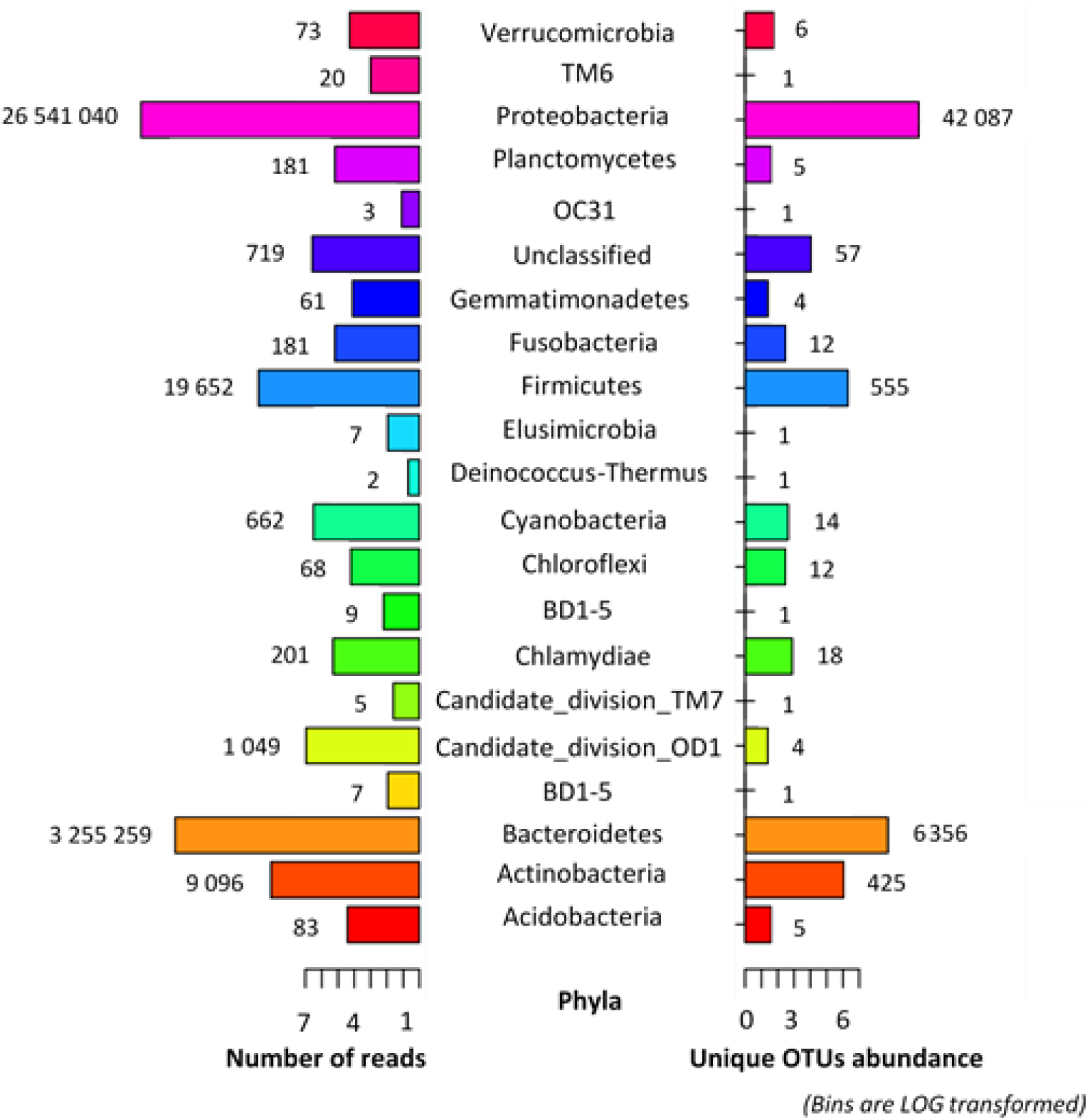
Number of reads per unique OTU abundance (at the phylum level). A comparison of the number of individual reads to the number of unique OTUs shows that phyla with high number of reads do not result in single OTUs.

**Figure S4.**
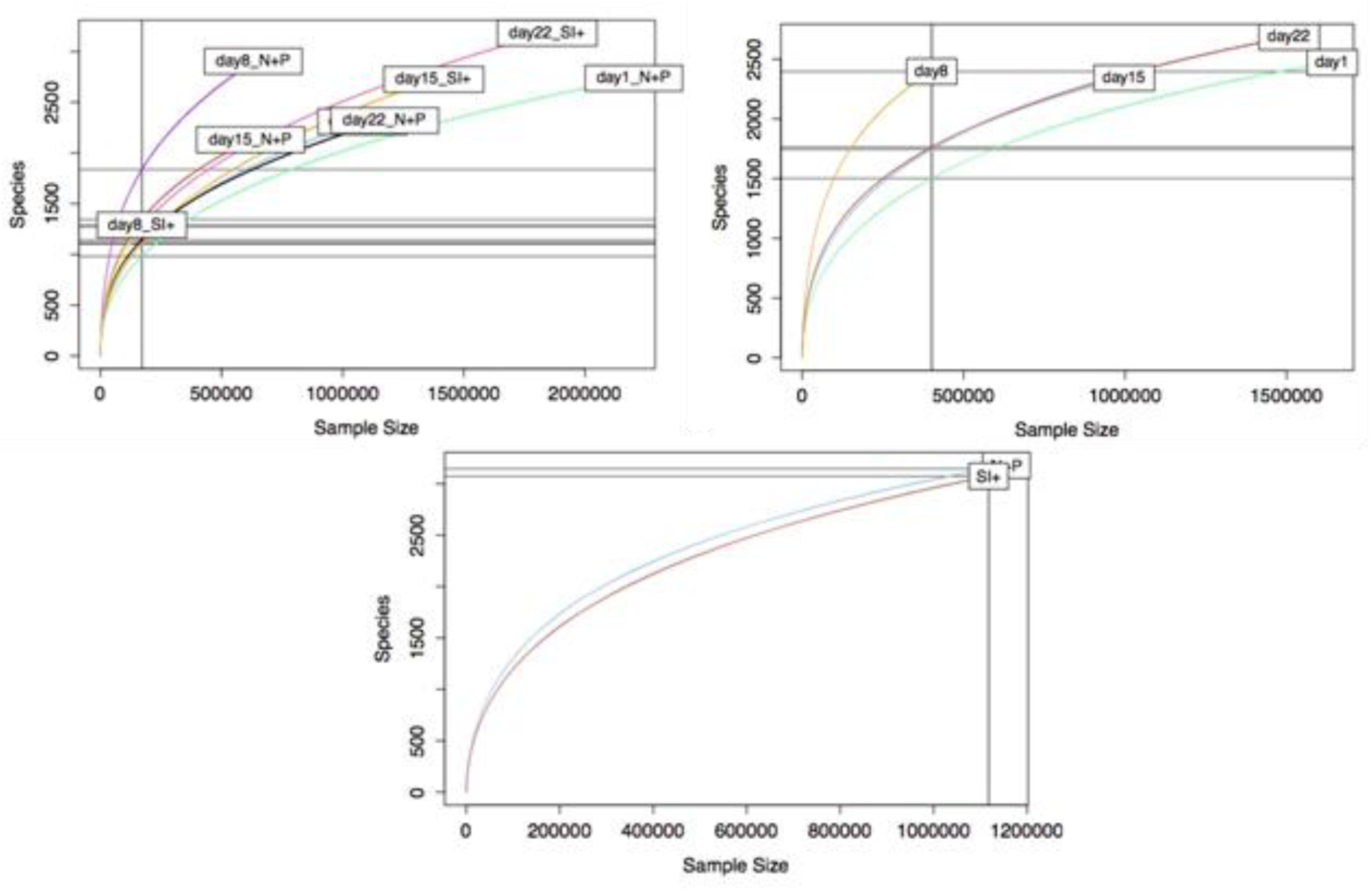
Alpha diversity. Rarefaction curves were used to evaluate the Alpha diversity in the different media conditions as well as at the different time points. Species richness in both minimal and complete media was ~3 000. Species richness over time remained between ~2 400 and 2 600, with reduced species richness (~1 300) on Day 8 (both minimal and complete media) possibly due to elevated levels of 16S *P. tricornutum* chloroplast reads which had to be omitted. Greatest species richness (~ 3 000) was shown on Day 22.

**Figure S5.**
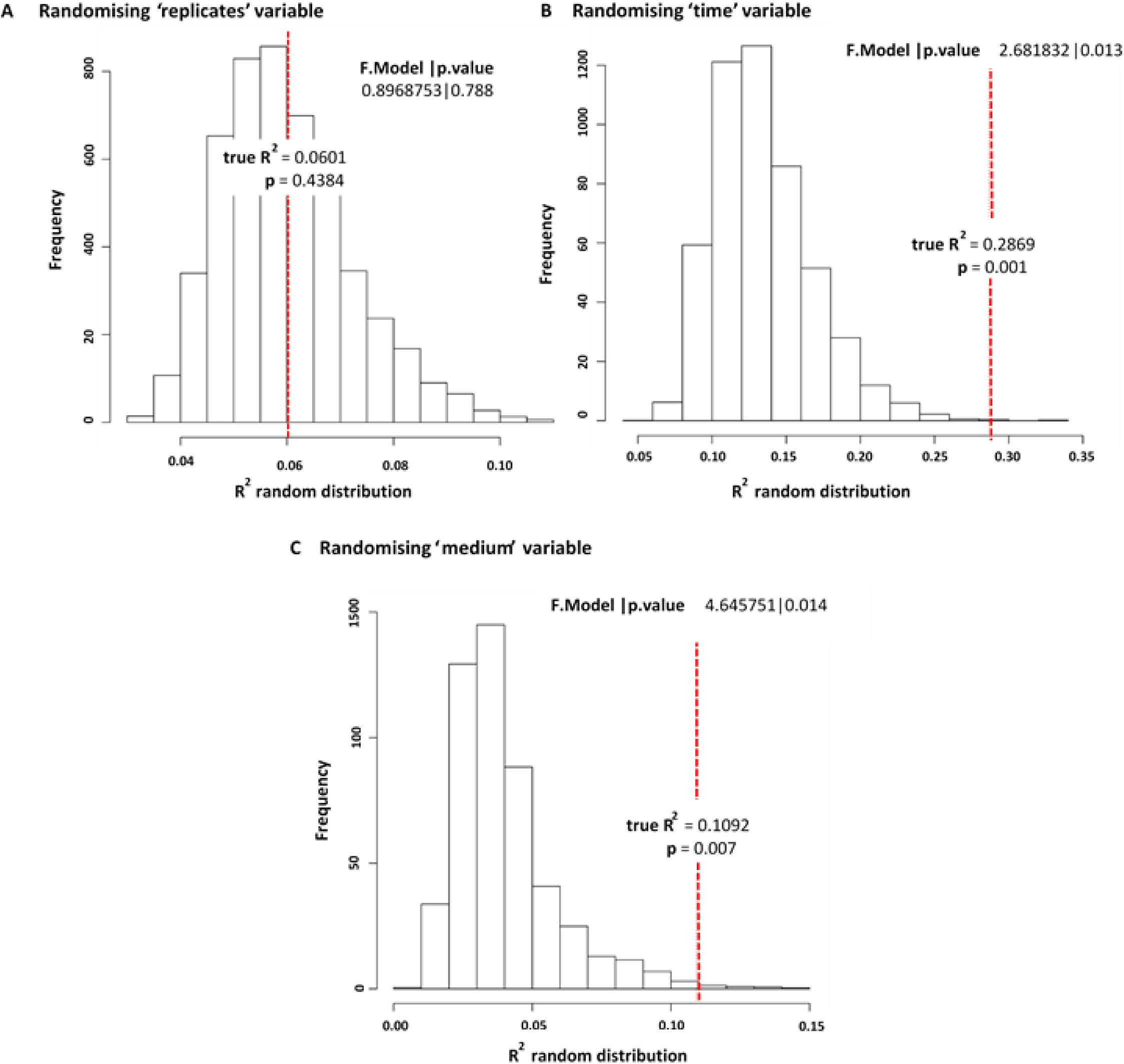
Beta diversity. A modified version of PermanovaG was used to carry out permutational multivariate analysis of variance using multiple distance matrices. The distance matrices [24x24] were previously calculated based on the generalised UniFrac distance (Chen *et al.*, 2012), weighted UniFrac and unweighted UniFrac (Lozupone and Knight, 2005) distance. The significance for the test was assessed by 5000 permutations. **A** shows no significant effect between the replicates (p-value of 0.4384). **B** shows a significant effect for the time variable (p-value of 0.001). **C** shows also shows a significant effect for the medium variable (p-value of 0.007)

**Figure S6.**
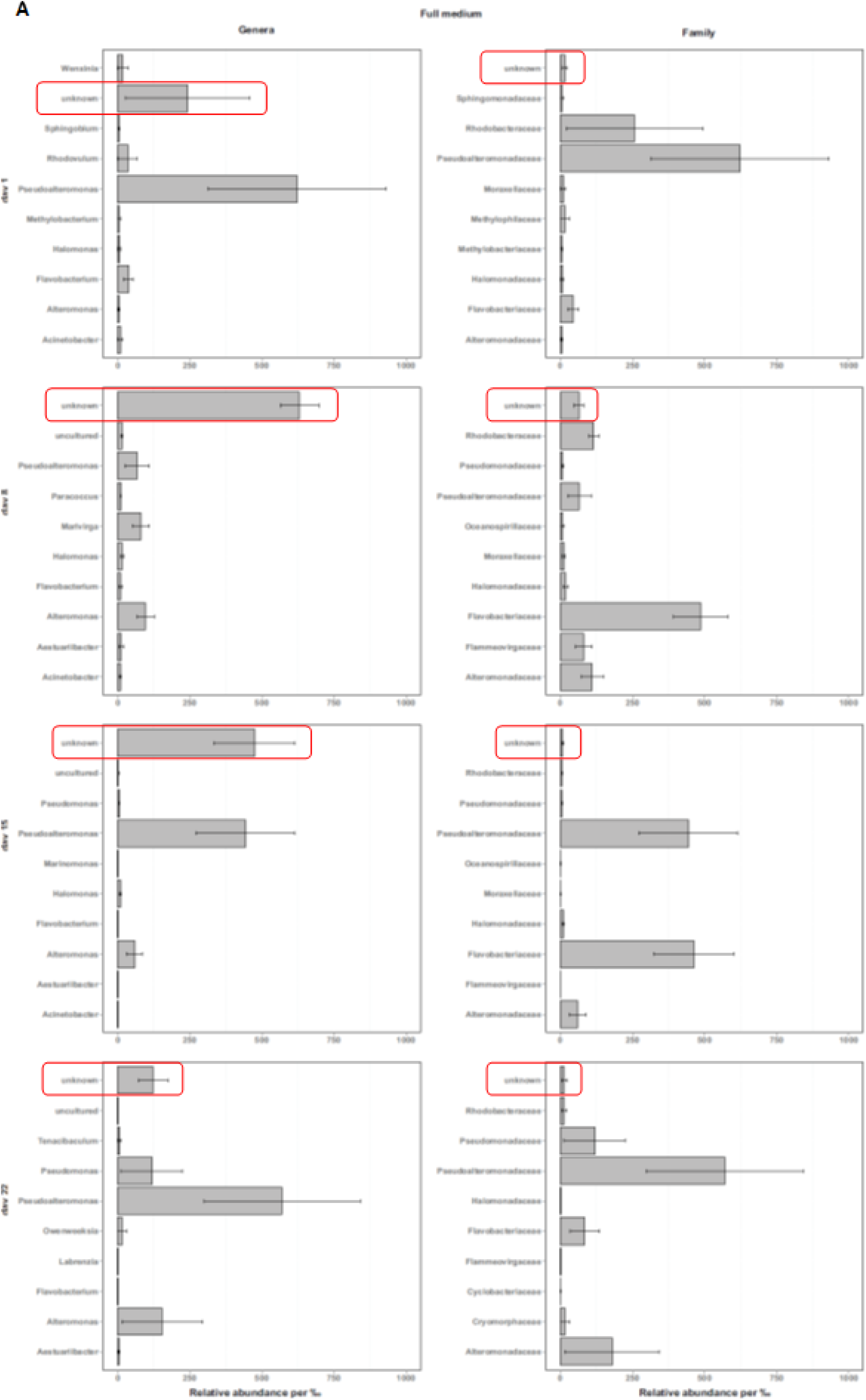
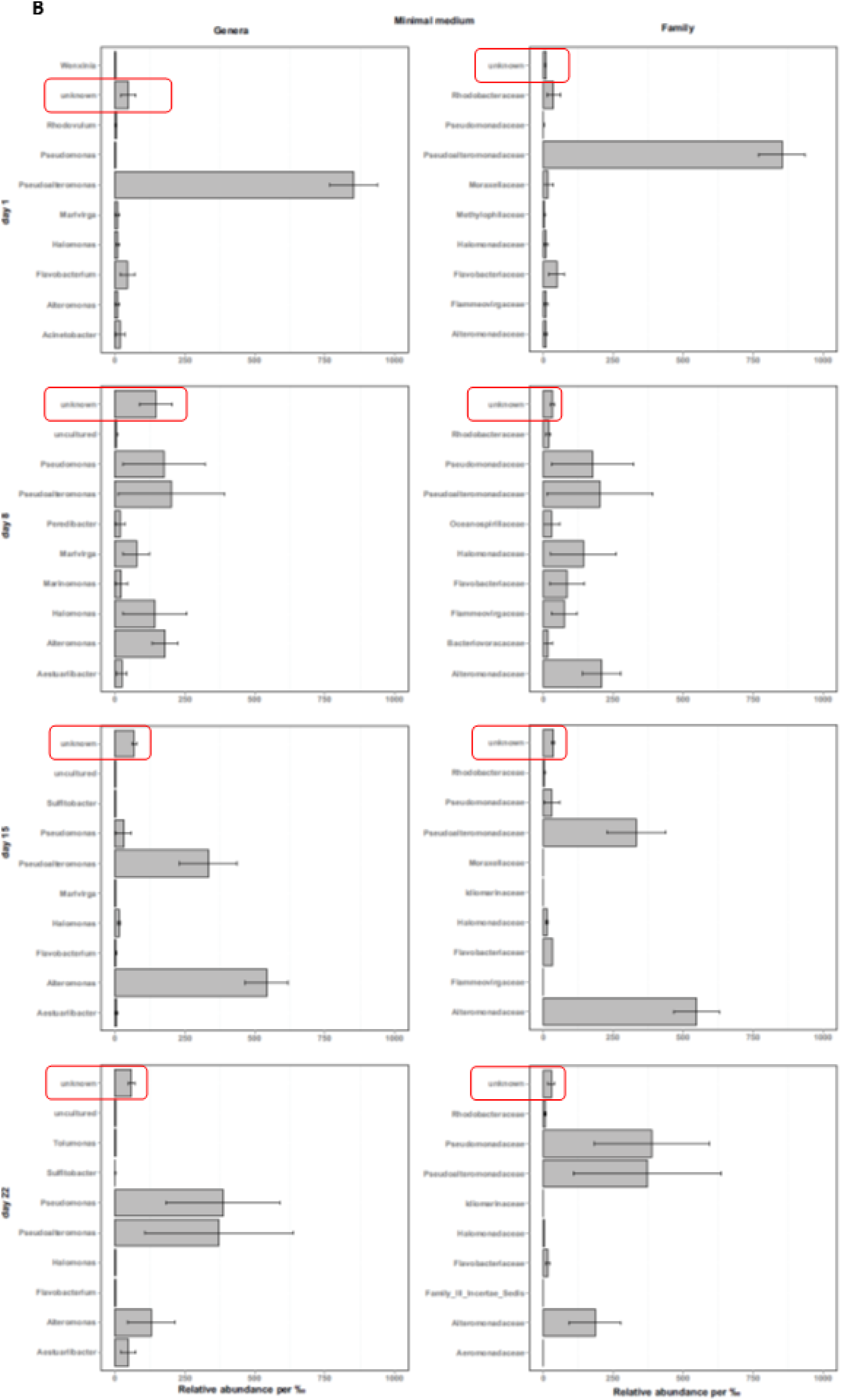
Comparison between bacterial community at genera level and family level. **A** in complete media. **B** in minimal media. We show no dynamical difference within the genera that cannot be observed at the family level. Encircled in red, there are a greater number of OTUs that could not be assigned a taxonomy (‘unknowns’) at the genera level than at the family level. By investigating the bacterial community dynamics at the family level, we also include taxonomical information that is unavailable at the genus level.

**Table S1.**
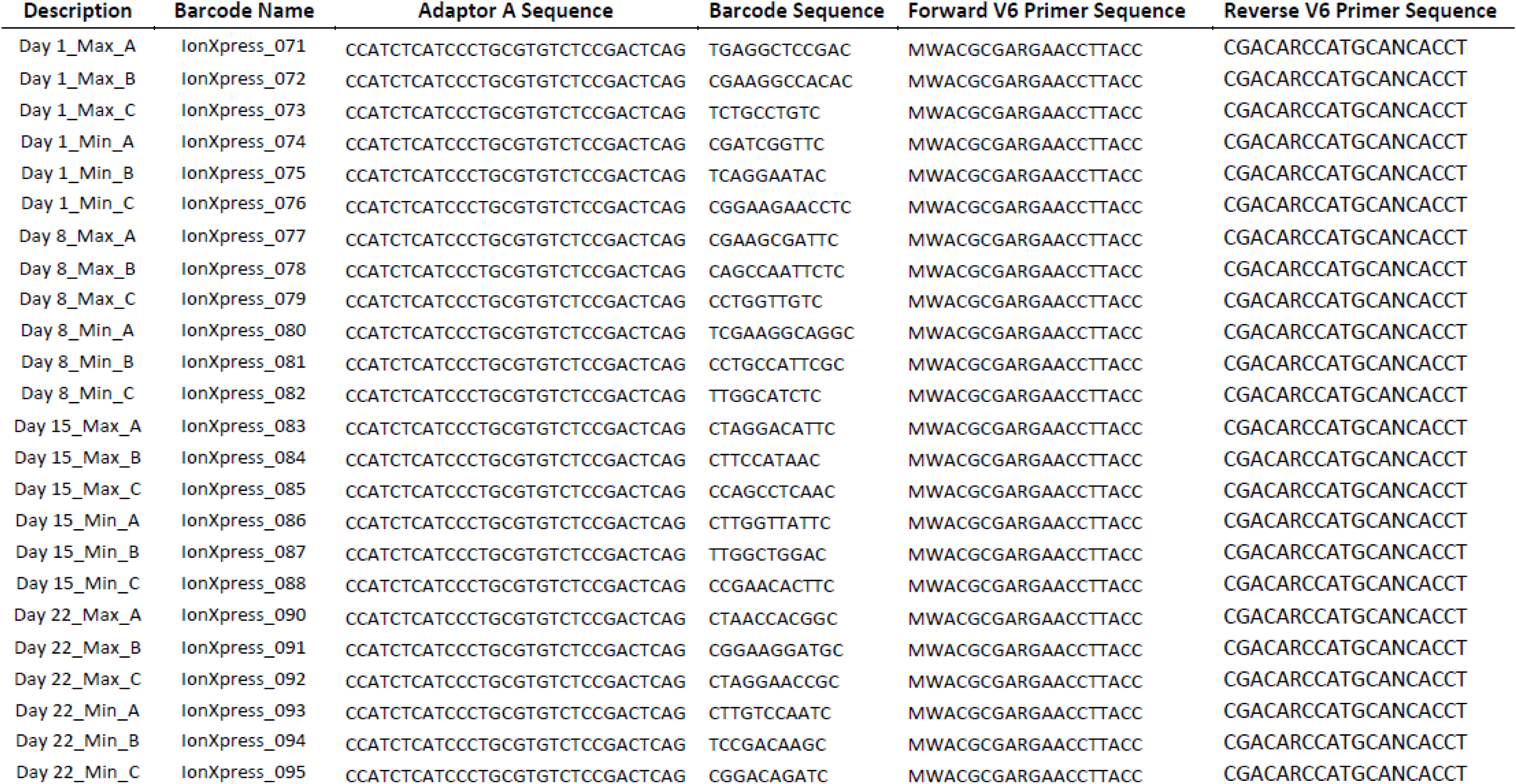
16S V6 rRNA primer sequences. ‘Max’ is the complete media. ‘Min’ is the minimal media. ‘A’, ‘B’, and ‘C’ are the three replicates.

**Table S2.**
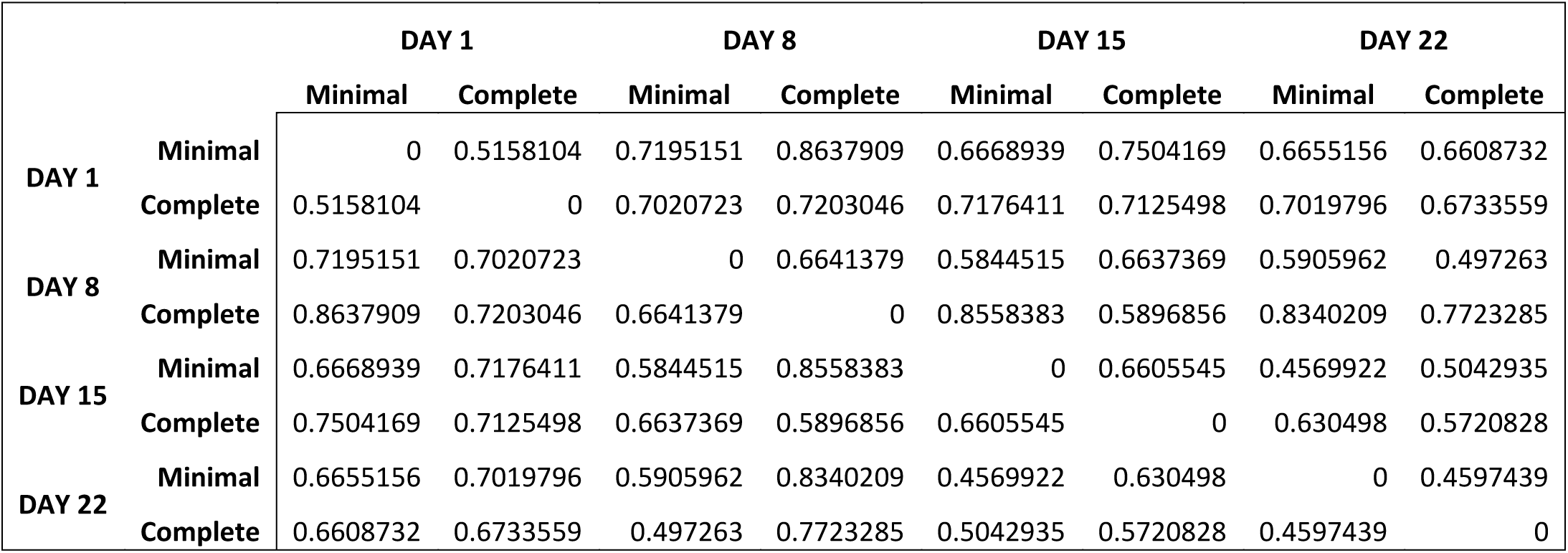
Generalised UniFrac distances of bacterial communities in complete and minimal media over time. Generalised UniFrac distance contains an extra parameter α controlling the weight on abundant lineages so the distance is not dominated by highly abundant lineages. α = 0.5 has overall the best power.

|  |  | DAY 1 |  | DAY 8 |  | DAY 15 |  | DAY 22 |  |
| --- | --- | --- | --- | --- | --- | --- | --- | --- | --- |
|  |  | Minimal | Complete | Minimal | Complete | Minimal | Complete | Minimal | Complete |
| DAY 1 | Minimal | 0 | 0.5158104 | 0.7195151 | 0.8637909 | 0.6668939 | 0.7504169 | 0.6655156 | 0.6608732 |
|  | Complete | 0.5158104 | 0 | 0.7020723 | 0.7203046 | 0.7176411 | 0.7125498 | 0.7019796 | 0.6733559 |
| DAY 8 | Minimal | 0.7195151 | 0.7020723 | 0 | 0.6641379 | 0.5844515 | 0.6637369 | 0.5905962 | 0.497263 |
|  | Complete | 0.8637909 | 0.7203046 | 0.6641379 | 0 | 0.8558383 | 0.5896856 | 0.8340209 | 0.7723285 |
| DAY 15 | Minimal | 0.6668939 | 0.7176411 | 0.5844515 | 0.8558383 | 0 | 0.6605545 | 0.4569922 | 0.5042935 |
|  | Complete | 0.7504169 | 0.7125498 | 0.6637369 | 0.5896856 | 0.6605545 | 0 | 0.630498 | 0.5720828 |
| DAY 22 | Minimal | 0.6655156 | 0.7019796 | 0.5905962 | 0.8340209 | 0.4569922 | 0.630498 | 0 | 0.4597439 |
|  | Complete | 0.6608732 | 0.6733559 | 0.497263 | 0.7723285 | 0.5042935 | 0.5720828 | 0.4597439 | 0 |

## Supplementary Model Material: Mathematical model of the population dynamics in *P. tricornutum* associated bacterial community

### 1. Model description

In order to test our understanding emerging from the analysis of the experimental data we built a Ordinary Differential Equations (ODE) model to simulate how the bacterial community develops in time. Considering the limited information that can be extracted from the experimental data, the model is purely qualitative and provides a proof of concept that a quantitative model can be constructed if dedicated experiments are designed for calibration. We base the organism-to-organism interactions on the production/consumption of metabolites. Metabolites include nutrients, micronutrients and toxins.

#### 1.1 Introduction

We develop a dynamic model represented by a set of 13 ODEs. Five ODEs describe the variation in time of the populations of *P. tricornutum (D)*, *Pseudoalteromonas (PA)*, *Flavobacterium (F)*, *Alteromonas (A)* and *Pseudomonas (P)*. The other eight ODEs describe the production and consumption of the metabolites we consider as mainly contributing to drive the community dynamics: the dissolved organic carbons of preference for *P A* and *A* (DOC_*P* A_ and DOC_*A*_, respectively), the complex polymers (COP) consumed by *F*, generic vitamins (Vit) and iron (Fe) needed by *D* and produced by *A*, bactericidial molecules (EPA and Bac, produced by *D* and by *P A* respectively) and the dissolved organic matter (DOM).

The model is built from the following working hypotheses:

1. the growth *γ* of each population follows a standard Verhulst equation parametrized with a carrying capacity *CC* and scaled by Michaelis-Menten-like terms that encode the dependence on necessary nutrients as a scaling factor 0 *< ʵ <* 1;
2. the death *δ* of each population is inversely proportional to (1+*γ*) to account for the fact that cells during replication (high growth rate) are healthier;
3. additional contributions to population death is given by the presence in the environment of noxious elements like bactericidal substances;
4. changes in metabolite concentrations are in general directly proportional to the growth *γ* of the consumers and producers;
5. for the DOC_*A*_ and COP metabolites we introduce the hypothesis that, in the event of micronutrient scarcicity, the diatom *D* will secrete more organic carbons favorited by those bacteria (*A* in our case) able to provide the needed micronutrients (Fe and Vit in our model);

Despite its simplicity and the minimal amount of assumptions made to build it, this model has 55 unknown free parameters (5 carrying capacities *CC*, 34 maximal rates *v*, 15 “Michaelis-Menten like” constants *K*, the fraction of DOC_*A*_-dependent growth *E*_DOC*A*_).

#### 1.2 Equations

Five ODEs describe the variation in time of the populations of organism *O*, with *γ^O^* and *δ^O^* being its growth and death rate:

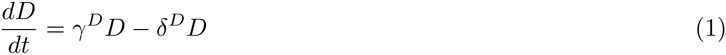

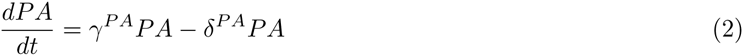

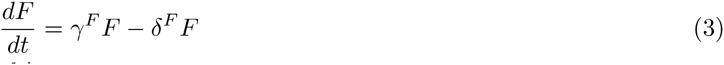

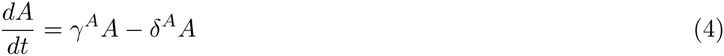

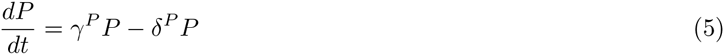

Eight ODEs describe the variation in time of the metabolites *J*, with 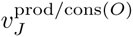 being the maximal production/consumption rate of *J* by organism *O*:

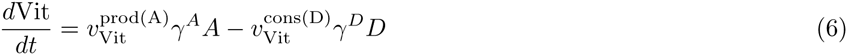

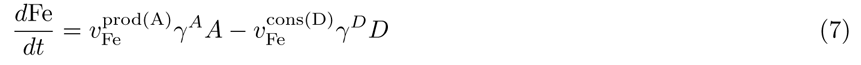

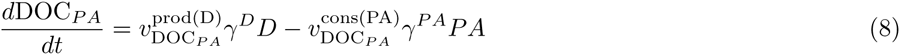

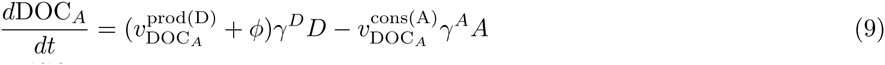

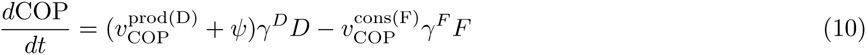

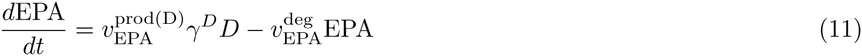

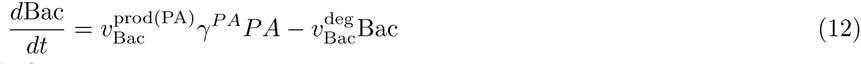

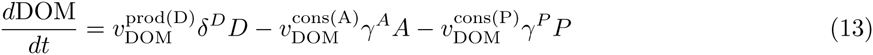

*φ* and *ψ* are additional terms for DOC_*A*_ and COP production respectively (see Section 1.3). 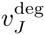 degradation rate of the bactericidal substances. Organism *O* growth and death rates depend in general on carrying capacity *CC^O^*, maximal rates 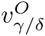 and on metabolites concentrations *J* with Michaelis-Menten-like constants 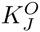 and eventually maximal rates 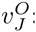:

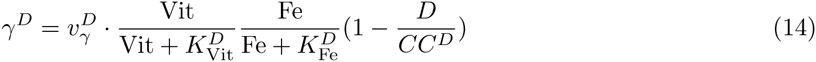

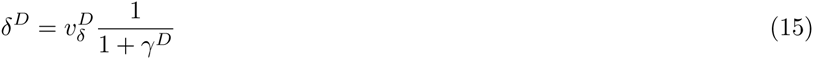

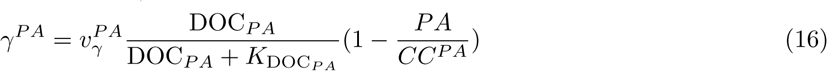

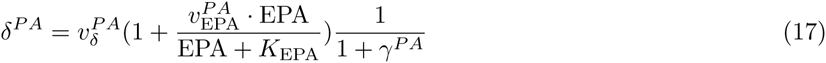

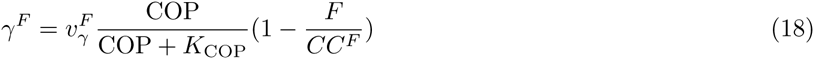

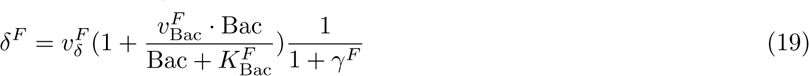

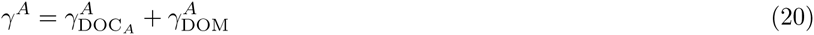

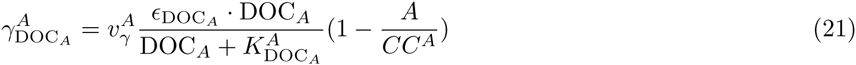

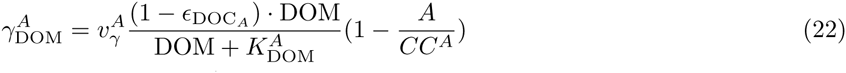

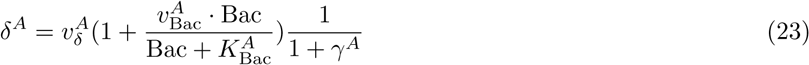

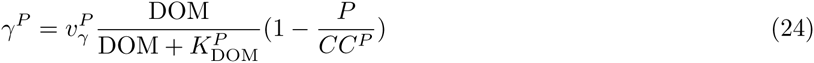

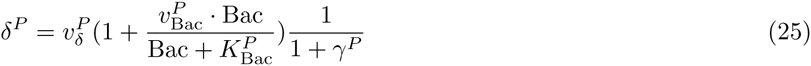

In the case of *A*, where growth is thought to be sustained by two different complementary nutrients, the final growth *γ* can be represented as the sum of two terms 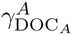 and 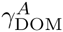 (Equations 21 and 22), with the parameter 0 < *E*_DOC*A*_ < 1.

#### 1.3 DOC_*A*_ and COP production

When *D* is grown in minimal media conditions, the emergence of *A* is observed over *F*. From this observation we hypothesise that *D* can produce extra organic carbons for either *A* or *F* depending on the scarcicity of micronutrients to favor the growth of *A* if more Vit or Fe is needed. We model the production of DOC_*A*_ and COP (Equations 9 and 10) introducing the functions *φ* and *ψ* defined as:

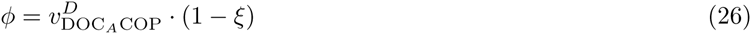

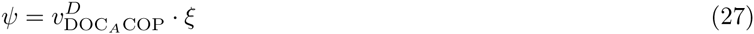

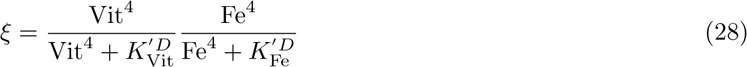

where 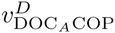 is the maximal additional production rate and 0 *< ξ <* 1 depends on Vit and Fe with fourth order Hill equations terms parametrised with 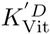 and 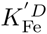 (see Figure 1).

**Figure 1:**
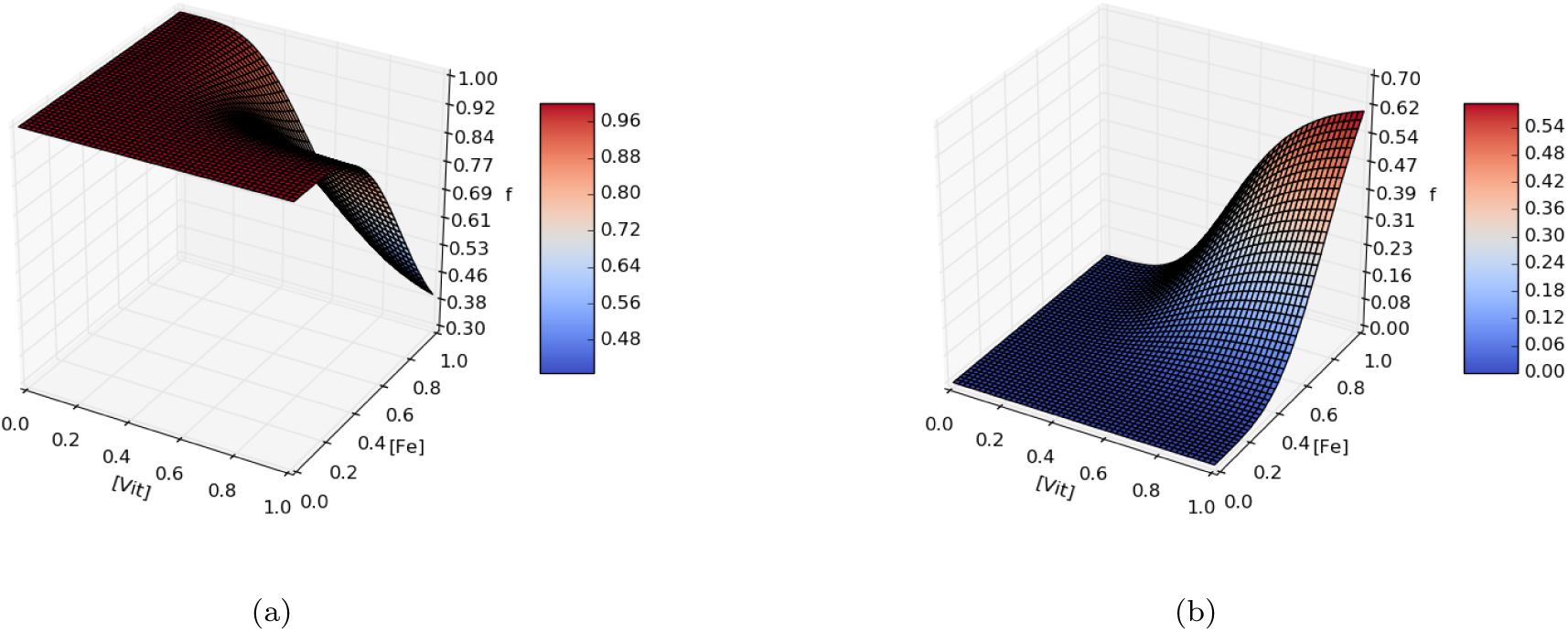
Example for DOC_*A*_ ((a), 1 − *ξ*) and COP ((b), *ξ*) additional production rates dependent on Vit and Fe availability in the media. Here 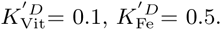 = 0.1, *K^lD^* = 0.5.

### 2 Parameters choice

The model has 56 parameters, of which 55 are free parameters (see Table 1). Being a qualitative model, we do not aim at interpreting the absolute parameter values in a biological sense. We can however draw considerations from relative values and stability tests.

**Table 1:** Total number of parameters for each parameter set. The dependent parameter is *ɛ*_DOM_= 1 *− E*_DOC__*A*_ in the sub-set of *A* parameters *P*(*A*).

| Parameter sub-set | $\mathcal{P}(D)$ | $\mathcal{P}(PA)$ | $\mathcal{P}(A)$ | $\mathcal{P}(F)$ | $\mathcal{P}(P)$ | degradation |
| --- | --- | --- | --- | --- | --- | --- |
| Sub-set size | 15 | 9 | 14 | 8 | 8 | 2 |

#### 2.1 Parameter fitting

The available data that can be used to fit the model parameters are the diatom biomass growth in two media conditions and four time points with bacteria relative abundances again in two media conditions. We can therefore fit the diatom biomass *D* evolution and the four relative bacteria *i* abundances *B*_*i*_/Σ_*j*_*B*_*j*_ time-course.

We implement as general strategy a genetic algorithm, where an “individual” *i* is a full set of 56 parameters 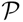, a “population” is an *ensemble* of parameter sets {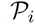}, a population at a certain evolution step is a “generation” and “evolution” goes as:

1. the first generation {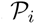{^0^ is populated by extracting the parameters from random uniform distributions within user-chosen ranges;
2. for each 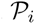 the ODE system is solved and a fitness score (see Section 2.1.1) is computed;
3. the most fit 10% individuals are retained as parents for the next generation;
4. the remaining individuals have a probability *P* = 0.05 to be also selected as parents;
5. parents are crossed to obtain enough children to reach the original population size;
6. crossing means randomly pick a parameter sub-set from one parent or the other;
7. each children has a probability *P* = 0.3 to randomly mutate one parameter;
8. the process is repeated from step 2. until generation 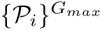.

##### 2.1.1 Fitness score

Fitness scores are computed in a different way when fitting the diatom growth or the bacteria relative abundances. When fitting to the diatom biomass data we compute the score as a simple euclidean distance:

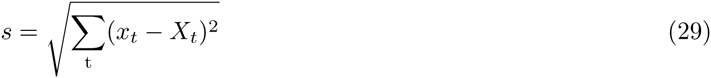

where the sum over time extends over 22 time points, *x*_t_ is the *D* biomass at time *t* and *X*_t_ is the biomass data at time *t*. The lower *s*, the better the fit. This score definition works well to fit the measurements of diatom biomass, but presents a big problem when used with bacteria relative abundances. A relative abundance is a number between 0 and 1, and we observe high variations including bacteria population going from very close to 0 to high abundance. Having only three time points to fit (the first 16S measurement is used as initial point), it can happen that constantly low abundant population are kept by the algorithm. We therefore define for the fit of bacteria relative abundances the following score:

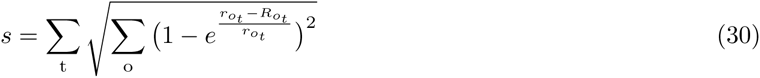

where the sum over time extends over 3 time points and the sum over organisms over the 4 bacterial species, *r*_*ot*_ is the relative abundance from the ODEs system solution for organism *o* at time *t* and *R*_ot_ is it the corresponding experimental relative abundance. This score definition allows to penalize the event of population extinction: when *r* is 0, the exponential term is 0 and the score is 1, while when *r* = *R* the exponential term is 1 and the score is 0 (see Figure **??**).

**Table 2:** Datasets used to fit diatom growth in minimal and complete media (MM and CM respectively). Time is scaled (1/3 of a day) to fit reasonably the growth phases (lag-log-exp-decay) using parameters 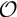(1). For the same reason cell counts are scaled to bring the lower count close to 0, but not feature-scaled to avoid loosing information on differences among MM and CM conditions. Only average values, and not experimental errors, are taken into account.

| T | 8 | 16 | 40 | 48 | 64 | 72 | 104 | 112 | 120 | 128 | 152 |
| --- | --- | --- | --- | --- | --- | --- | --- | --- | --- | --- | --- |
| MM | 0.004 | 0.021 | 0.133 | 0.325 | 0.820 | 1.012 | 1.121 | 1.187 | 1.192 | 1.233 | 1.209 |
| CM | 0.050 | 0.044 | 0.162 | 0.605 | 0.733 | 0.919 | 1.037 | 1.099 | 1.134 | 1.108 | 0.859 |
| T | 168 | 176 | 184 | 208 | 216 | 232 | 240 | 248 | 264 | 272 | 288 |
| MM | 1.104 | 1.096 | 0.951 | 1.015 | 0.965 | 0.851 | 0.869 | 0.704 | 0.481 | 0.504 | 0.394 |
| CM | 0.821 | 0.844 | 0.624 | 0.682 | 0.624 | 0.556 | 0.535 | 0.478 | 0.199 | 0.282 | 0.303 |

**Table 3:** Relative abundances of the four bacterial families at three intermediate time points (days 8, 15 and 22). The abundances were scaled from the experimental values (where more families were present) to add to unity.

| t | Complete Media |  |  |  | Minimal Media |  |  |  |
| --- | --- | --- | --- | --- | --- | --- | --- | --- |
|  | PA | F | A | P | PA | F | A | P |
| 64 | 0.101 | 0.724 | 0.159 | 0.014 | 0.294 | 0.132 | 0.308 | 0.264 |
| 120 | 0.453 | 0.474 | 0.061 | 0.010 | 0.351 | 0.031 | 0.585 | 0.031 |
| 176 | 0.600 | 0.084 | 0.189 | 0.126 | 0.385 | 0.020 | 0.187 | 0.406 |

##### 2.1.2 Results of the genetic algorithm

The chosen population size is 200 and the algorithm stops either after non significant increase in fitness or at generation number 50. The algorithm can be run to fit six scenarios:

- D-MM: *D* Biomass in Minimal Media;
- D-CM: *D* Biomass in Complete Media;
- B-MM: Bacteria relative abundances in Minimal Media;
- B-CM: Bacteria relative abundances in Complete Media;
- D*B-MM: *D* Biomass and Bacteria relative abundances in Minimal Media;
- D*B-CM: *D* Biomass and Bacteria relative abundances in Complete Media;

For D-type fits, the fitness score of Eq. 29 is used. For B-type fits, the fitness score of Eq. 30 is used. For D*B-type fits, the fitness score is the product of the two scores. We will refer to D-fit, B-fit and D*B-fit in the following if media is not to be specified.

Considering the fact that a simple ODE model cannot capture metabolic readjustment, we do not expect to obtain the same parameters for CM and MM conditions. The fitting is therefore performed separately in the two conditions and in the following steps:

1. B-fit is run 20 times varying all 55 parameters in *O*(1) ranges
2. The parameters from the best B-fits are kept (*P_MM_*_1_ and *P_CM_*_1_)
3. After checking the effect of varying the different parameters sets (see Figure 2 and Section **??**), different variation ranges are chosen to perform refits
4. D*B-CM is run 5 times varying *P*(*D, deg*)_*CM*1_ *±* 50%, *P*(*A, F, P*)_*CM*1_ *±* 20%, *P*(*P A*)_*CM*1_ *±* 10%
5. D*B-MM is run 5 times varying *P_MM_*_1_ *±* 50%, and the best parameters are kept (*P_MM_*_2_)
6. D*B-MM is run again 5 times varying *P*(*D*)_*MM*2_ *±* 5%, *P*(*A, F, P, P A, deg*)_*MM*2_ *±* 80%

Stability analysis (see Figure 2 and Section **??**) shows that the only parameters from other sub-sets influencing the biomass growth curve in CM are 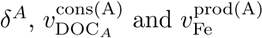, but we ignore them for these first iterations.

**Figure 2:**
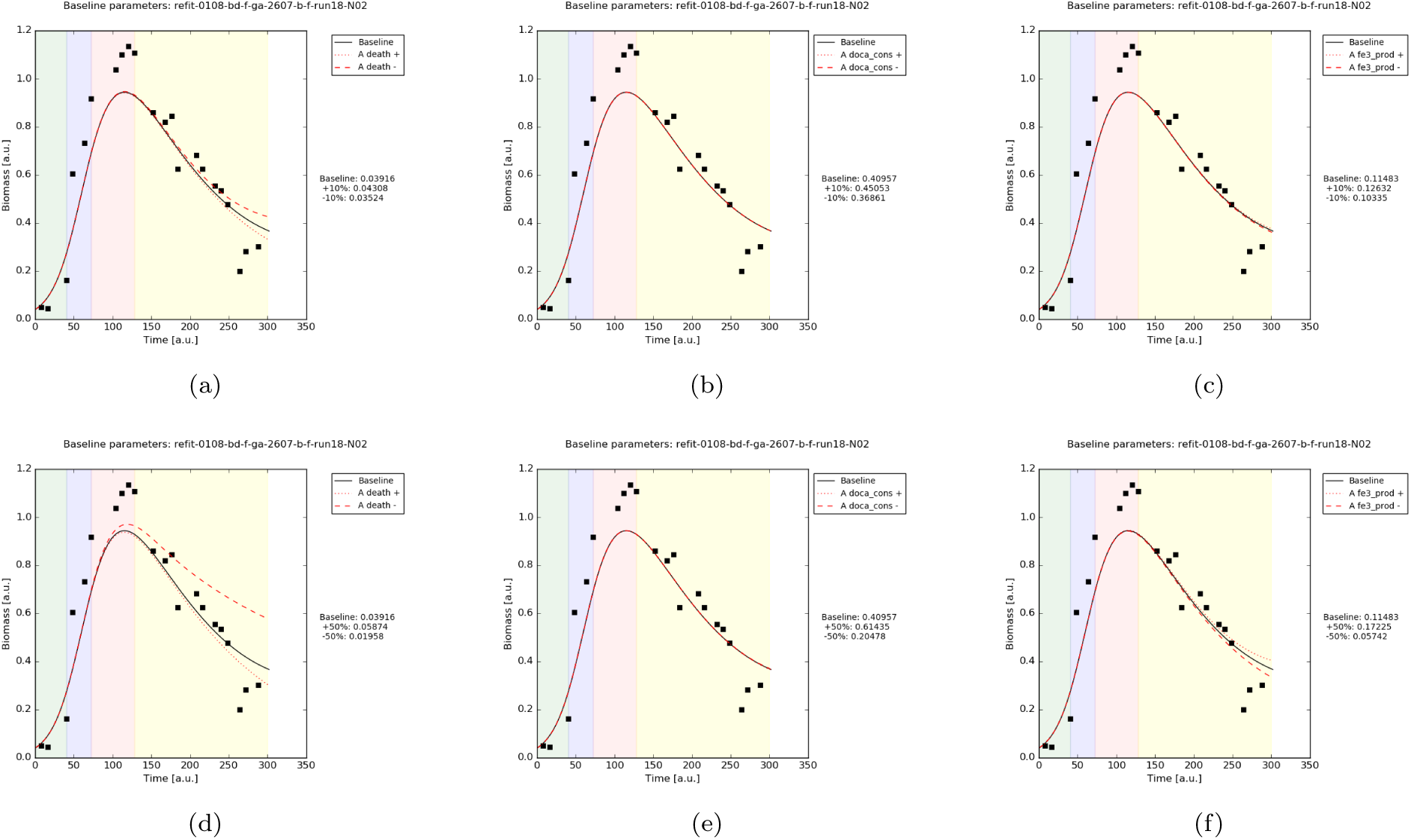
Diatom growth in MM simulation results. The parameters 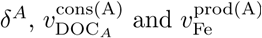, are varied by *±*10% (a, b, c respectively) and by ±50% (d, e, f respectively).

**Figure 3:**
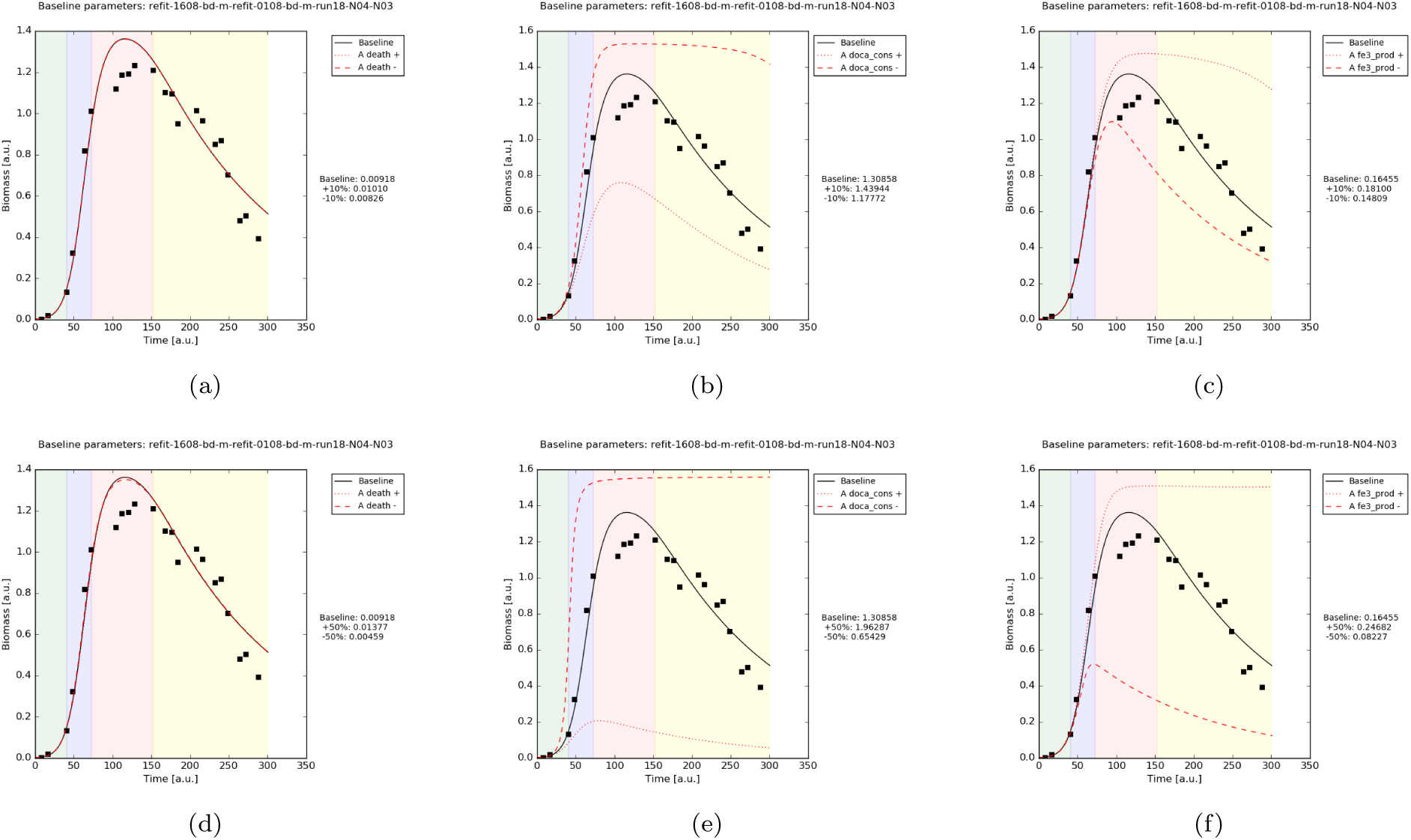
Diatom growth in CM simulation results. The parameters 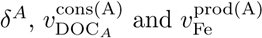, are varied by *±*10% (a, b, c respectively) and by ±50% (d, e, f respectively).

